# Exonuclease ISG20 inhibits human cytomegalovirus replication by inducing an innate immune defense signature

**DOI:** 10.1101/2025.02.10.637376

**Authors:** Matthias Hehl, Myriam Scherer, Eva-Maria Raubuch, Claudia Ploil, Teresa Rummel, Philipp Kirchner, Nina Kottmann, Anna Reichel, Anna Katharina Kuderna, Anne-Charlotte König, Caroline C. Friedel, Florian Erhard, Thomas Stamminger

## Abstract

ISG20 is an interferon-regulated protein that exhibits RNase activity thereby inhibiting the replication of a broad spectrum of RNA viruses. By single cell RNA sequencing, we identified ISG20 as an antiviral factor for human cytomegalovirus (HCMV) as it was upregulated in a population of HCMV-resistant cells. In accordance with an antiviral role on herpesviruses, overexpression of ISG20 in primary human fibroblasts led to reduced HCMV and HSV-1 replication, while knockdown of ISG20 enhanced virus growth. In Western blot kinetics, we observed that inhibition of HCMV replication by ISG20 occurs at the early stage of infection which correlated with reduced amounts of viral early and late transcripts. However, neither the half-life of viral and cellular RNAs nor of viral DNA was decreased in ISG20-expressing cells, indicating that ISG20 does not exert its antiviral effect via a degradation of RNAs or DNA. Instead, RNA-seq analysis revealed an innate immune defense signature upon ISG20 expression that comprised the upregulation of a distinct set of interferon stimulated genes (ISGs), zinc finger proteins (ZNFs) and of transposable elements (TEs). Our data indicate that this gene signature augments both IFN production and response of the host cell. Consistently, the JAK-STAT inhibitor ruxolitinib rescued HCMV gene expression in ISG20-expressing cells. We conclude that ISG20 induces a so far unprecedented immune defense signature that serves to amplify the IFN-mediated host cell defense thus explaining its broad antiviral activity.

## Introduction

The type I interferon (IFN) system plays a critical role in the innate immune defense, enabling rapid detection of invading pathogens and inducing an antiviral state in infected and neighboring cells. Viral nucleic acids thereby present pathogen-associated molecular patterns (PAMPs) to cellular pattern recognition receptors (PRRs), including DNA sensors like cyclic GMP-AMP synthase (cGAS) and RNA sensors such as retinoic acid-inducible gene-I (RIG-I) [1]. Once activated, these sensors recruit adaptor proteins and trigger intracellular signaling cascades that ultimately lead to the synthesis of type I IFNs, including IFN-α and IFN-β. The secreted IFNs bind to the IFN-α/β receptor (IFNAR) thereby activating Janus kinase 1 (JAK1) and tyrosine kinase 2 (TYK2) and leading to the recruitment of signal transducer and activator of transcription 1 and 2 (STAT1 and STAT2). Together with interferon-regulated factor 9 (IRF9), STAT1 and STAT2 assemble into the interferon-stimulated gene factor 3 (ISGF3) complex. Upon nuclear translocation, the ISGF3 complex binds to interferon-stimulated response elements (ISREs) in the promoter regions of interferon-stimulated genes (ISGs), inducing the production of hundreds of antiviral proteins [2].

The interferon-stimulated gene 20 (ISG20), a member of the death effector domain-containing (DEDD) exonuclease superfamily, exhibits an extended antiviral activity and has been linked to tumorigenesis [3–5]. First identified in 1997 as a PML nuclear body-associated and interferon-induced protein, ISG20 is primarily activated by type I interferons (IFNs) [6]. Rather than through ISGF3, ISG20 expression is stimulated by IRF1, which itself is induced by IFNs and engages an interferon-stimulated response element (ISRE) in the ISG20 promoter region [7]. Additionally, activation of the NF-κB pathway by double-stranded RNA (dsRNA) has been shown to rapidly induce ISG20 expression via a κB element in its promoter [8].

ISG20 is known to act as a 3’ to 5’ exonuclease harboring three distinct exonuclease domains (Exo I, II, and III) and is able to degrade single-stranded RNA (ssRNA) and DNA (ssDNA) *in vitro*, with a strong preference for RNA [9]. Due to this property, the antiviral role of ISG20 has been extensively studied and demonstrated against a wide range of RNA viruses, including positive-strand RNA viruses (hepatitis C virus, Zika virus, West Nile virus, dengue virus, hepatitis A virus, human Immunodeficiency virus type 1) and negative-strand RNA viruses (Vesicular Stomatitis Virus, Influenza A virus) [10–13]. However, some RNA viruses appear to be resistant, like severe acute respiratory syndrome coronavirus type 1, or inhibited only in certain cell-types, such as yellow fever virus [14]. Concerning DNA viruses, only limited data is available, with antiviral effects of ISG20 mainly characterized in the context of hepatitis B virus [15, 16].

Several reports describe that the antiviral function of ISG20 relies on its exonuclease activity, which is severely impaired by an aspartate-to-glycine mutation (D94G) in the Exo II domain of ISG20 [9–11]. Therefore, a direct degradation of viral nucleic acids by ISG20 has been postulated as the primary mechanism of viral inhibition [10]. Degradation of viral RNA by ISG20 is well-documented for hepatitis B virus (HBV), where ISG20 targets the ε-stem-loop structure within the HBV pregenomic RNA (pgRNA) through its Exo III domain, leading to accelerated decay of HBV pgRNA [15–17]. Interestingly, a recent study proposed that ISG20 also affects the cccDNA (covalently closed circular DNA) of HBV, in cooperation with the interferon-stimulated protein APOBEC3A. In this model, APOBEC3A deaminates cytosines to uracils on transcriptionally active cccDNA, which results in an accumulation of ISG20 and reduction of the HBV DNA [18]. Contrary to these findings, however, a number of studies observed viral inhibition in the absence of nucleic acid degradation. One study attributed ISG20’s antiviral effect against alphaviruses (Chikungunya virus) to the upregulation of ISGs in murine fibroblasts (MEFs) rather than to RNA degradation [19]. The upregulated proteins, especially IFIT1, thereby mediate inhibition of viral translation, which was confirmed in a mouse model with ISG20-deficient mice [19]. Further, ISG20’s antiviral effect against pseudorabies virus in pig epithelial cells was linked to IFN-β stimulation and consequent increased expression of ISGs and cytokines [14]. Increased IFN-β production was also observed in ovarian cancer cells with elevated ISG20 expression, which was suggested to result from an ISG20-induced accumulation of short dsRNA fragments activating the RIG-I signaling pathway [20]. In accordance with the inhibition of alphavirus translation, a translation inhibition specific for viral (non-self origin) but not cellular (self origin) RNAs has been demonstrated in the context of vesicular stomatitis virus (VSV) infection [21]. However, the authors neither detected an induction of IFN or IFIT1 by ISG20 expression in epithelial HEK293T and myeloid U937 cells, nor an involvement of IFIT1 in translation inhibition. In conclusion, while the antiviral function of ISG20 is well established, the underlying molecular mechanisms leading to viral inhibition appears to be subject to virus-or/and cell-type specific differences that remain to be further defined.

Here, we identify ISG20 as an antiviral factor against the DNA virus human cytomegalovirus (HCMV), a ubiquitous β-herpesvirus that can cause serious disease in immunocompromised individuals. As with all herpesviruses, lytic replication of HCMV is characterized by three consecutive phases of gene expression, the immediate early (IE), early (E), and late (L) phase. Using ISG20 overexpression and knockdown approaches, we found that ISG20 negatively influences the E and L phases of replication, dependent on an intact exonuclease domain. While this antiviral activity could not be attributed to the degradation of viral RNA or viral DNA, we observed that ISG20 enhances the expression of other ISGs during HCMV infection.

Characterization of ISG20-induced transcriptomic changes by ISG20 using RNA-seq and SLAM-seq analysis revealed an induction of not only ISGs but also a broad upregulation of transposable elements (TEs) and zinc-finger proteins in primary human fibroblasts. Thus, we describe a so far unprecedented immune defense signature upon ISG20 expression that is supposed to be critical for inhibition of human cytomegalovirus replication.

## Results

### Identification of ISG20 as antiviral factor against HCMV

Since previous studies have demonstrated a high cell-to-cell variability during viral infection, we performed single cell RNA sequencing (scRNA-seq) in order to identify host factors that modulate the early phases of lytic HCMV infection [22–24]. For this, primary human foreskin fibroblast (HFF) cells were infected with HCMV (strain AD169) at a multiplicity of infection of 0.4 and harvested at 6 hours post-infection (hpi) for droplet-based single-cell sequencing. The analyzed cells clustered mainly based on cell cycle markers, the amount of viral RNA, and the degree of interferon (IFN) activation (Figure 1A-C). Initiation of lytic infection led to formation of cluster 4, containing cells with high levels of viral immediate-early and early RNAs. Notably, cells in cluster 3, which displayed low levels of immediate-early transcripts UL122/123 and no detectable transcripts with slower expression kinetics, such as UL37, showed a clear antiviral signature [25] (Figure 1B, C). The marker genes of this cluster included several factors with well-established antiviral functions, such as the interferon-stimulated genes (ISGs) *IFIT1/2/3* [26] (Figure 1C).

**Figure 1.**
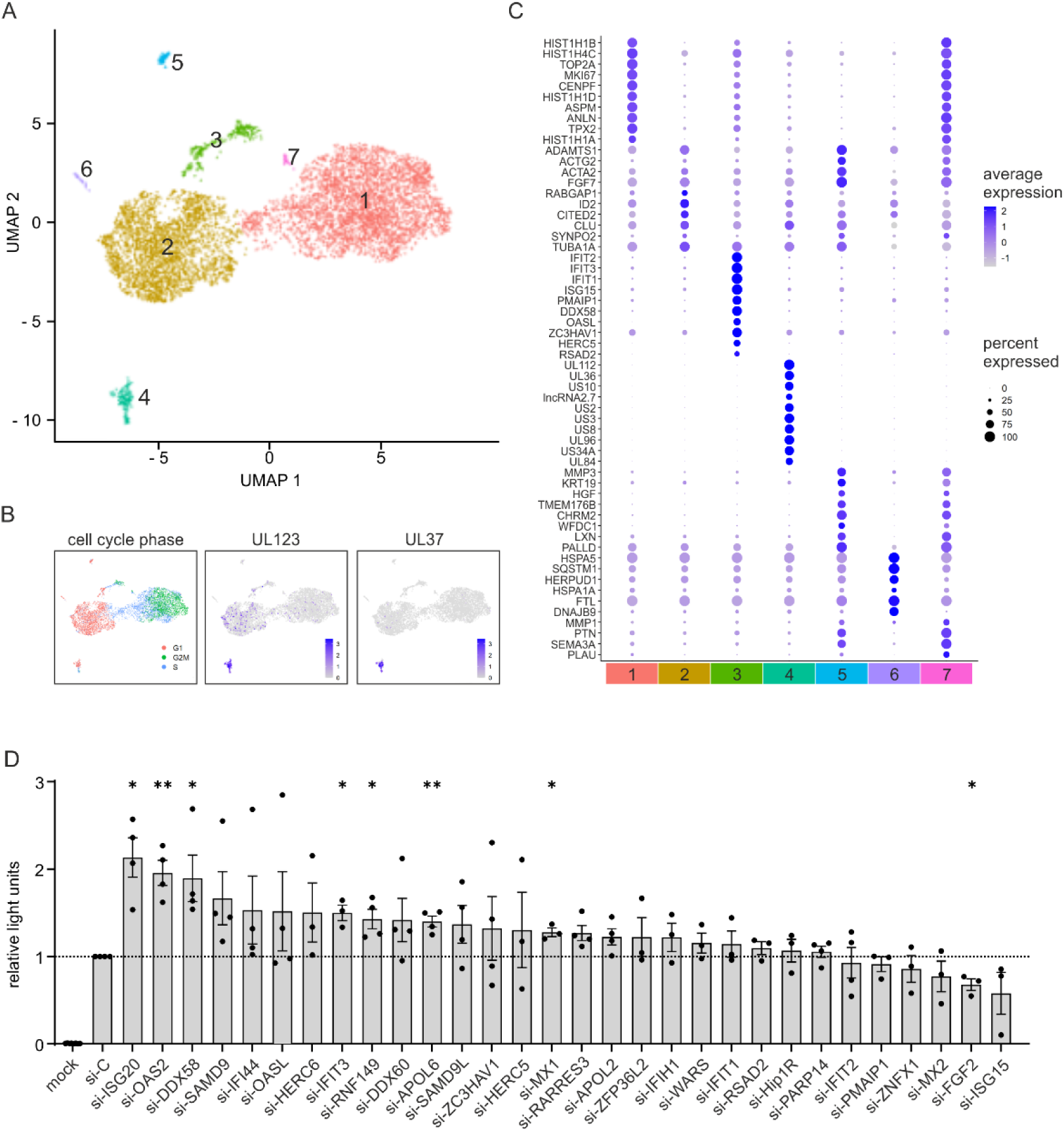
Identification of ISG20 as antiviral factor restricting HCMV replication. (A-C) Single cell RNA-sequencing of HFF cells infected with HCMV (strain AD169, MOI 0.4) for 6 h. (A) UMAP projection of infected HFF cells color-coded according to supervised clustering of gene signatures. (B) UMAP plots with cells colored by cell cycle phase (left panel), expression of HCMV UL123 (middle panel) and HCMV UL37 (right panel). (C) Dot plot representing expression of marker genes in the HFF cell clusters. Circle diameter represents the percentage of cells expressing a particular gene, normalized average expression is represented by color intensity. (D) SiRNA screen of potential antiviral genes identified by single cell RNA-sequencing. HFF cells were transfected with control siRNA (si-C) or siRNAs against indicated genes using a custom siRNA-library (Qiagen). Two days later, cells were left non-infected (mock) or were infected with HCMV (strain TB40/E) expressing firefly-luciferase under control of the UL84 early promoter (MOI of 0.01). 7 days post-infection, viral replication was measured using a Luc-Screen luciferase assay (Applied Biosystems). Values are shown as relative mean values +/- SEM derived from at least three independent experiments performed in triplicates. Statistical analysis was performed using a one-sample t-test. *, p<0.05; **, p<0.01; no asterisk: not significant.

We next focused on the top 30 genes of this cluster and investigated whether they act as antiviral factors during HCMV infection. For this purpose, we knocked down the individual genes in HFF cells using a custom siRNA library, infected the cells with a luciferase-expressing HCMV and quantified viral replication at 7 days post-infection (dpi) using a luciferase assay. The majority of the 30 tested factors displayed an antiviral activity in this assay, although a significant impact on viral replication was only observed for 7 genes (Figure 1D). The most pronounced increase in luciferase expression was detected after depletion of *ISG20*, followed by *OAS2*, *DDX58*, *IFIT3, RNF149*, *APOL6*, and *MX1*. For *FGF2*, a significant proviral effect was observed, as its depletion led to a reduction in HCMV replication. Analysis of the knockdown efficiency by western blotting revealed an almost complete knockdown of *ISG20* (Supplementary Figure 1A), whereas a partial depletion was observed for the other analyzed factors *SAMD1*, *ZAP* and *TRIM28* (Supplementary Figure 1B). Furthermore, no major effects of siRNA transfection on cell viability were observed, as determined by measuring intracellular ATP levels in a Cell-TiterGlo assay (Supplementary Figure 1C, D). In summary, we identified several factors that inhibit HCMV infection, with the strongest effect detected for the interferon-stimulated gene *ISG20*.

### ISG20 is transiently upregulated during HCMV infection

Since ISG20 has hardly been characterized in the context of herpesvirus infections, we wished to further elucidate its role during lytic HCMV replication. Consistent with previous data, expression of the ISG20 protein could be detected in HFF cells after stimulation with interferons (IFNs), in particular with IFN-β, but not in unstimulated/uninfected cells [6] (Figure 2A). Western blot analysis of HCMV-infected cells revealed an upregulation of ISG20 early during infection, followed by a decrease in ISG20 protein levels, which appeared to be more pronounced in cells infected with HCMV strain TB40/E compared to AD169 (Figure 2B). This observation is in line with an initial induction of ISG expression caused by viral attachment and entry, and subsequent transcription inhibition by viral proteins, such as the immediate-early protein 1 (IE1). In fact, infection with an IE1-deleted HCMV induced a stronger and more permanent increase of ISG20 expression (Figure 2C). The changes of ISG20 protein levels during HCMV infection also correlated with changes in mRNA levels, as determined through RT-qPCR. Comparison of temporal profiles suggested a slightly delayed inhibition of ISG20 transcription in comparison to other ISGs and in particular to IFN-β, whose transcription was shut down more rapidly (Figure 2D). In summary, the regulation of ISG20 during HCMV infection resembles that of other ISGs with protein levels peaking during the early phase of infection.

**Figure 2.**
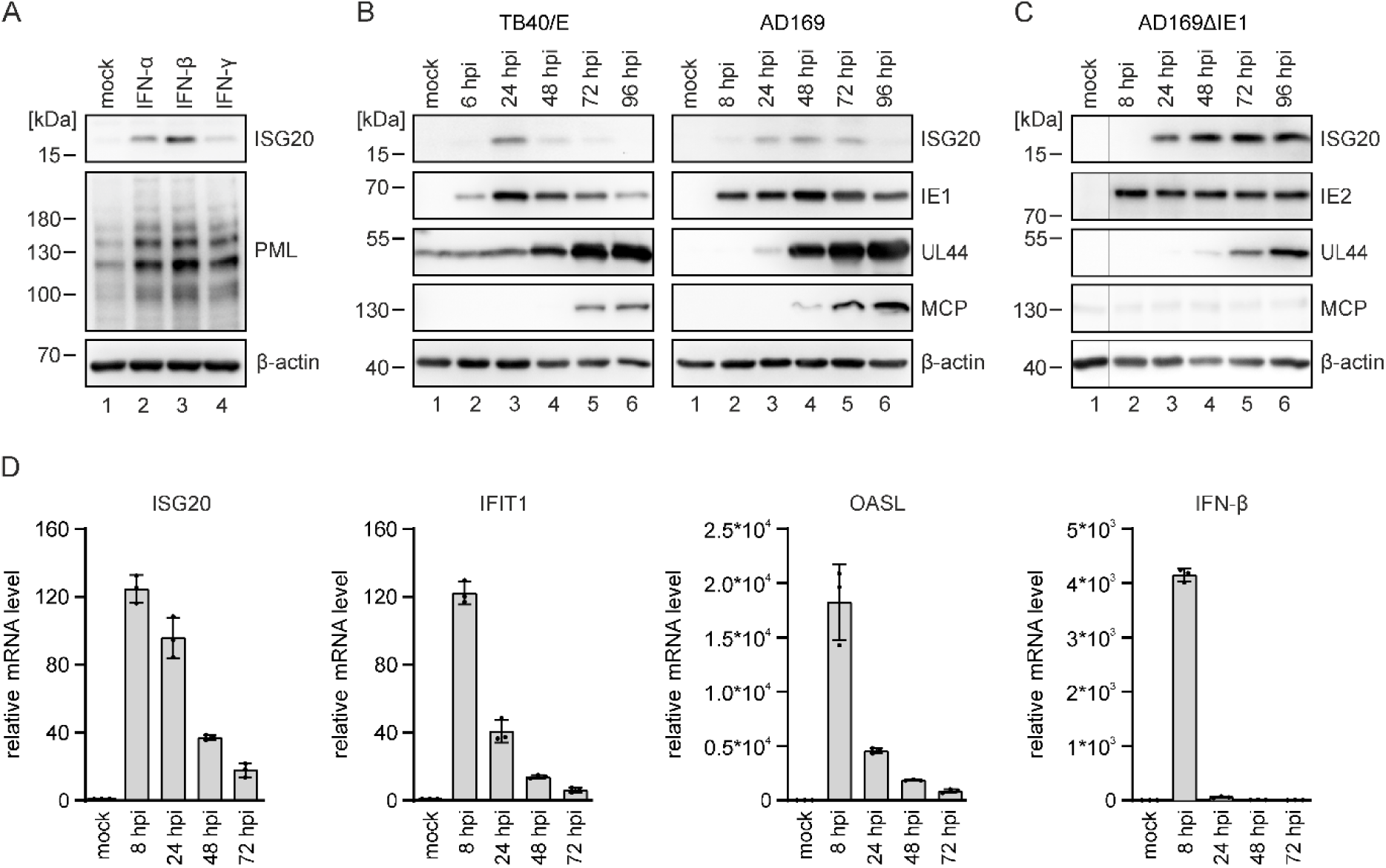
ISG20 expression during HCMV infection. (A) Induction of ISG20 expression by IFNs. HFF cells were left untreated (mock) or were treated with IFN-α (1000 U/ml), IFN-β (1000 U/ml) or IFN-γ (0.2 mg/ml). After 24 h, cells were harvested for western blot analysis of ISG20 expression. The interferon-inducible protein PML was included as positive control and β-actin as loading control. (B, C) ISG20 protein levels during HCMV infection. HFF cells were either not infected (mock) or infected with HCMV strain TB40/E at a MOI of 2.5 (B), AD169 at a MOI of 3 (B), AD169ΔIE1 at a MOI of 4 (C). At indicated times after virus inoculation, cell lysates were prepared and the expression levels of ISG20 as well as viral immediate early (IE1/IE2), early (UL44), and late (MCP) proteins were analyzed by western blotting. Cellular β-actin was detected as loading control. (D) ISG and IFN mRNA levels during HCMV infection. HFF cells were infected with HCMV strain TB40/E at a MOI of 3 or were not infected (mock). At indicated times after virus inoculation, cells were harvested and subjected to total RNA isolation and RT-qPCR to determine mRNA levels of ISG20, IFIT1, OASL and IFN-β. Values were normalized to the reference gene GAPDH and are shown as relative mean values derived from triplicates +/-SD. One representative experiment out of two is shown. hpi, hours post-infection.

### ISG20 overexpression limits HCMV and HSV-1 replication

To confirm the antiviral function of ISG20 during HCMV infection, we utilized lentiviral vectors to generate HFF cells with doxycycline-inducible overexpression of either wild-type ISG20 (HFF/ISG20) or the catalytically inactive D94G mutant (HFF/ISG20mut), alongside control cells (HFF/control). The expression of ISG20 and ISG20mut following doxycycline treatment was confirmed by western blotting (Supplementary Figure 2A). Since high expression levels of ISG20 have previously been associated with cellular toxicity, cell viability was assessed at different times after doxycycline stimulation [21]. In our inducible system, no toxic effects were observed, even after prolonged ISG20 expression over several days (Supplementary Figure 2B).

To characterize the effect of ISG20 overexpression on HCMV replication in the newly generated cells, multistep growth curve analysis was performed. HFF cells expressing ISG20 or the catalytically inactive ISG20 mutant as well as control HFFs were infected with HCMV strains TB40/E or AD169 (Figure 3A-C). Samples of cell supernatants were collected at various times after infection, and the number of viral genomes was determined by quantitative real-time PCR (qPCR). We observed a pronounced antiviral effect of ISG20, with both HCMV strains showing similarly reduced virus growth of about 1 log in ISG20-expressing cells compared to control cells. Notably, expression of catalytically inactive ISG20 caused a slight reduction in HCMV replication at a MOI of 0.1 (Figure 3A, B), while no effect of ISG20mut was detected upon infection with a lower virus dose (Figure 3C). Overexpression of ISG20, but not the exonuclease-dead mutant, also led to a potent inhibition of the α-herpesvirus HSV-1, as assessed by growth curve analysis using a VP22-GFP-encoding recombinant virus. These data suggest an antiviral function of ISG20 against members of the alpha- and beta-herpesvirus family that is dependent on its exonuclease activity.

**Figure 3.**
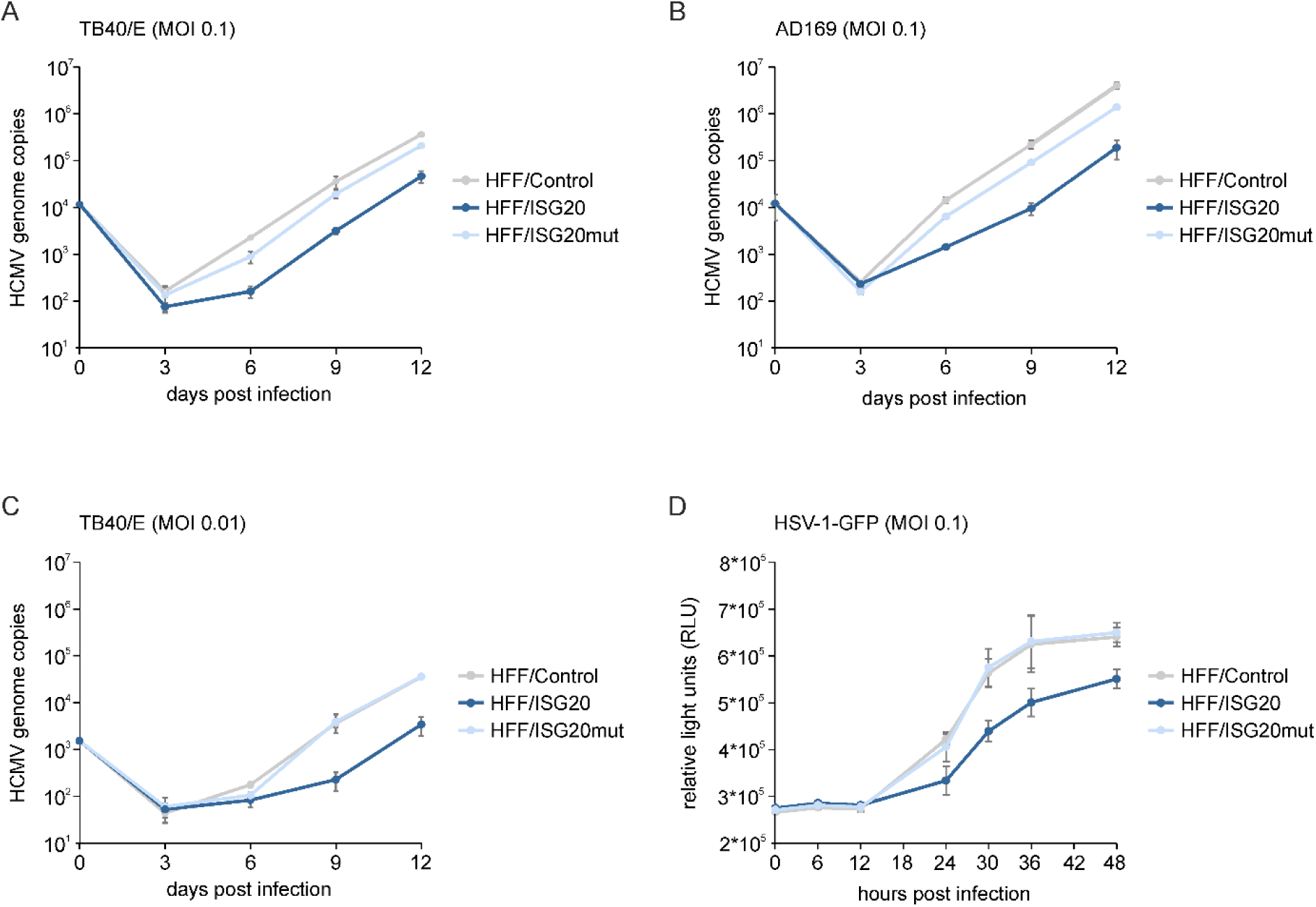
Effect of ISG20 overexpression on HCMV and HSV-1 replication. (A-C) Multistep growth curves of HCMV in HFFs with inducible ISG20 expression. HFF/Control, HFF/ISG20 and HFF/ISG20mut were seeded in triplicates and stimulated with doxycycline (500 ng/ml) for 24 h, before they were infected with HCMV strain TB40/E at a MOI of 0.1 (A) or 0.01 (C) or AD169 at a MOI of 0.1 (B). Cell supernatants were harvested at indicated times after infection and analyzed for genome copy numbers using an HCMV IE1-specific quantitative real-time PCR. Mean values +/-SD are shown. (D) Multistep growth curve of HSV-1 in HFFs with inducible ISG20 expression. HFF/Control, HFF/ISG20 and HFF/ISG20mut were seeded in triplicates and stimulated with doxycycline (500 ng/ml) for 24 h, before they were infected with VP22-GFP-expression HSV-1 at a MOI of 0.1. At specified times post-infection, the GFP-signal was measured in living cells using a luminescence microplate reader. Mean values +/-SD derived from three independent experiments are shown as relative light units (RLU).

### ISG20 negatively affects the early and late phase of HCMV replication

To elucidate which step of the HCMV replication cycle is affected by ISG20, viral protein accumulation was monitored during one round of replication. To this end, HFFs expressing ISG20 or ISG20mut as well as control HFFs were infected with AD169 or TB40/E (MOI of 1). At different times after infection, total cell lysates were prepared and subjected to western blotting to analyze the expression kinetics of viral immediate early (IE), early (E), and late (L) proteins. Interestingly, no negative effect of ISG20 on the viral immediate-early phase was observed (Figure 4A, B). Instead, we noted a slight increase in the expression of IE1 and IE2 in HFF/ISG20, which was corroborated by densitometric quantification (Figure 4C). Viral protein expression in the early and late phase, in contrast, was markedly decreased in HFF/ISG20 compared to HFF/Control, with similar effects observed for AD169 and TB40/E (Figure 4A-C). Expression of ISG20mut caused a mild inhibition of viral early/late protein accumulation. These observations were confirmed in a different set of cells with stable expression of ISG20-mGFP or ISG20mut-mGFP (Supplementary Figure 3). Again, a reduction of viral early and late protein levels was detected in ISG20-mGFP-expressing cells, albeit the effect was mainly visible under low MOI conditions which may be due to low expression levels of ISG20-mGFP after long-term cell passaging (Supplementary Figure 3B). To further substantiate our findings, viral protein accumulation was monitored in ISG20-depleted HFF. Transfection of HFF cells with an ISG20-specific siRNA prior to HCMV infection (strain TB40/E, MOI of 1) resulted in elevated levels of viral early and late proteins (Figure 4D), which appeared to be more pronounced when lower virus doses were applied (Figure 4E). In summary, these results suggest that ISG20 inhibits HCMV by negatively modulating the early and late phases of the viral replication cycle.

**Figure 4.**
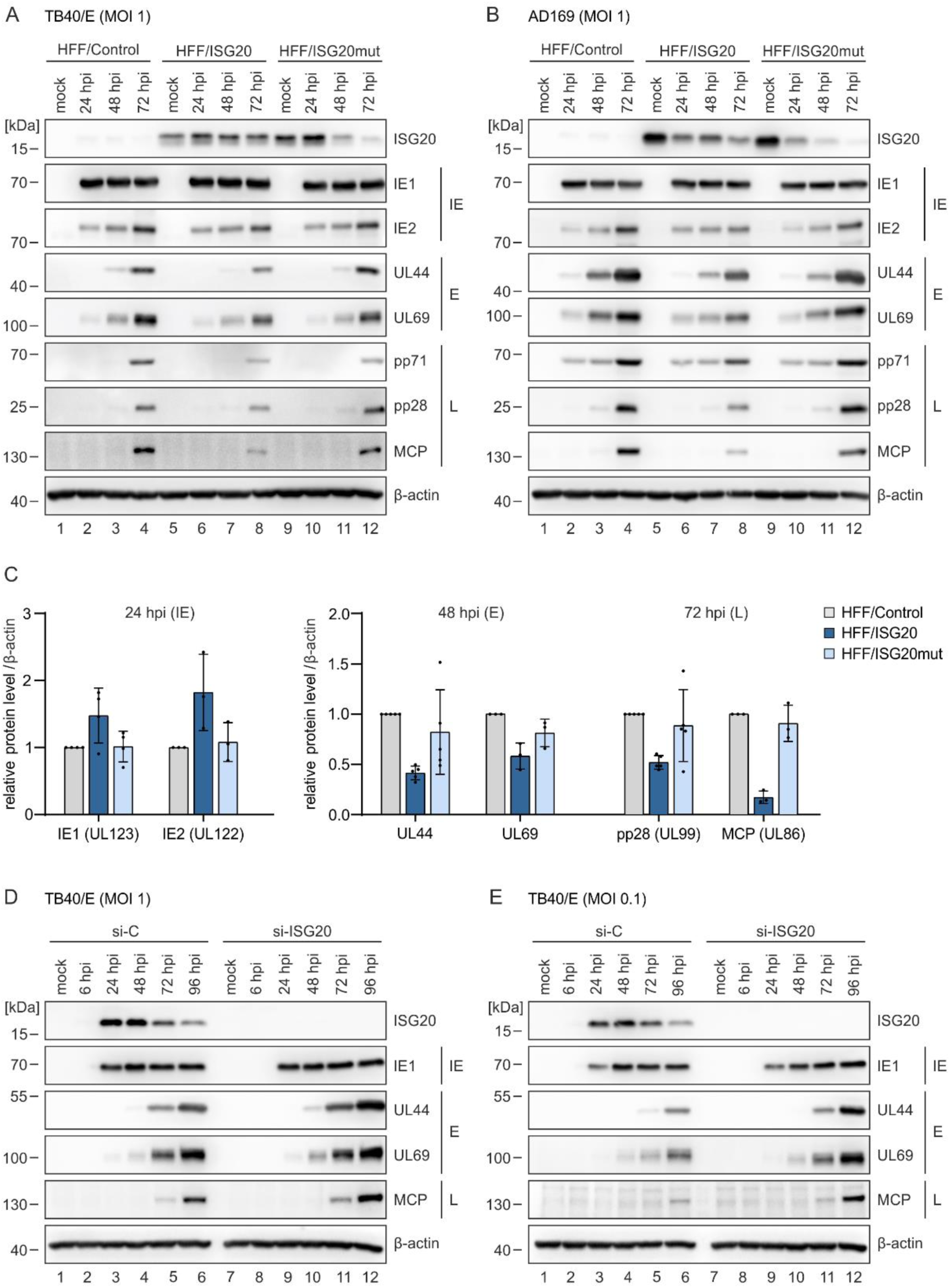
Effect of ISG20 on the expression pattern of HCMV proteins. (A-C) Influence of ISG20 overexpression on viral protein accumulation. HFF/Control, HFF/ISG20 and HFF/ISG20mut were stimulated with doxycycline (500 ng/ml) for 24 h, before they were infected with HCMV strain TB40/E (A) or AD169 (B, C) at a MOI of 1 or were not infected (mock). At indicated times after virus inoculation, cell lysates were prepared and the expression levels of viral immediate early (IE), early (E), and late (L) proteins were analyzed by western blotting. Cellular β-actin levels served as loading control. (C) Densitometric analysis of western blots was performed using ImageJ. Levels of viral proteins were normalized to β-actin and are shown as relative mean values +/- SD derived from ≥ 3 experiments, which were performed with independently transduced cell populations. (D, E) Influence of ISG20 knockdown on viral protein accumulation. HFF cells were transfected with an ISG20-specific or a control siRNA and, 3 days post-transfection, were infected with HCMV strain TB40/E at a MOI of 1 (D) or 0.1 (E). At indicated times after virus inoculation, cell lysates were prepared and the expression levels of viral IE, E, and L proteins were compared by western blotting. Cellular β-actin was detected as loading control. One representative experiment of two is shown. hpi, hours post-infection.

### ISG20 does not induce the degradation of HCMV DNA

Since the exonuclease ISG20 has been reported to induce the degradation of HBV cccDNA, we wondered whether the inhibition of HCMV infection by ISG20 may be attributed to the degradation of HCMV DNA [18]. To elucidate the effect of ISG20 on viral DNA, we first determined the amount of HCMV input genomes in HFF/ISG20 compared to HFF/ISG20mut and HFF/Control. Total DNA was extracted early during HCMV infection, before synthesis of new HCMV genomes takes place, and viral DNA copy numbers were quantified by qPCR. We detected similar genome numbers in all cell populations at 6 hpi, and a slight, not significant reduction of HCMV genomes in HFF/ISG20 compared to HFF/Control at 24 hpi, but no clear degradation of viral input DNA by ISG20 (Figure 5A). Comparable results were obtained in HFF transfected with ISG20-specific siRNA or control siRNA prior to HCMV infection. Overall, these data indicate that ISG20 does not inhibit HCMV replication via the degradation of incoming viral genomes (Figure 5B).

**Figure 5.**
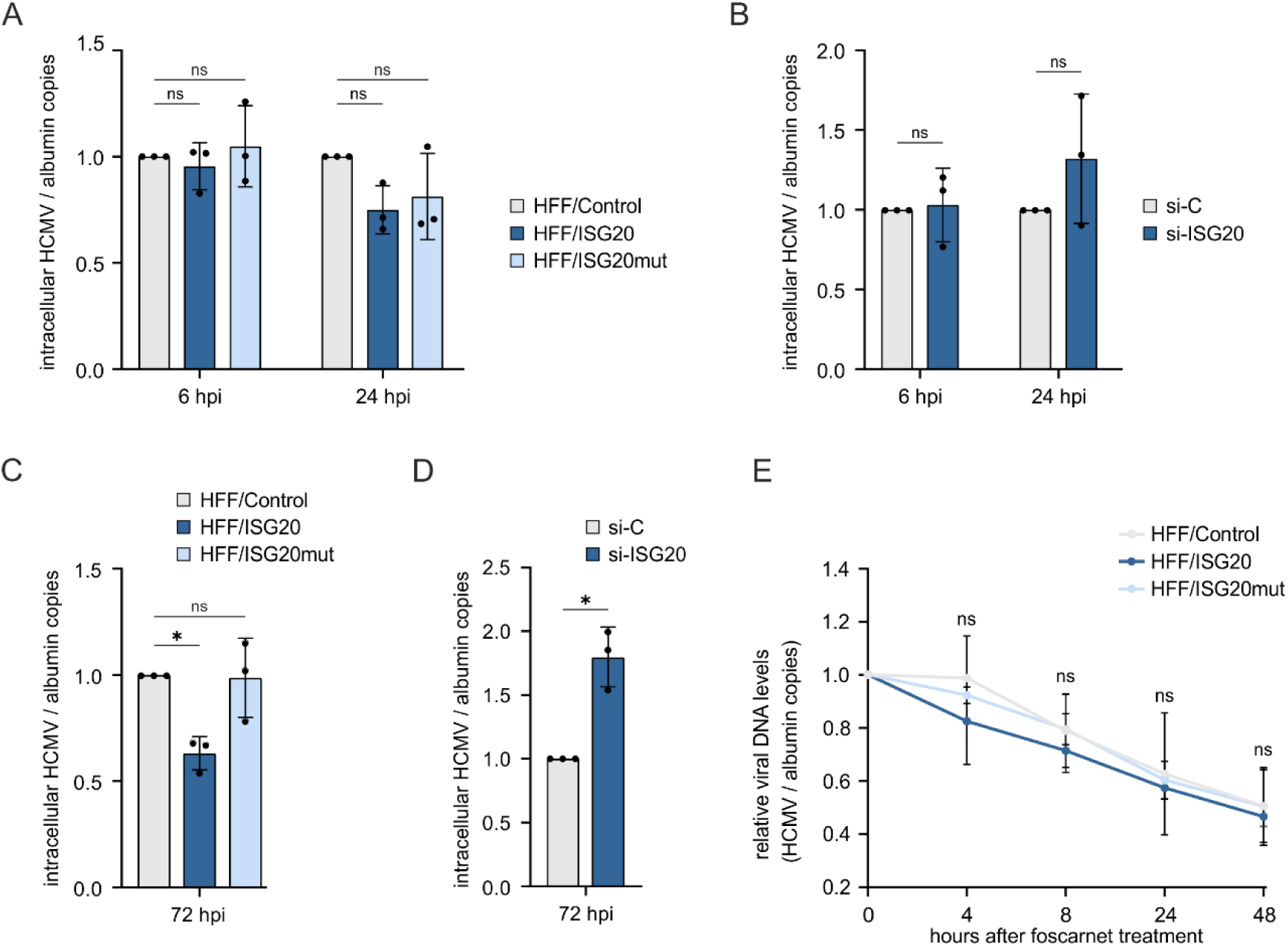
Effect of ISG20 on the stability of HCMV DNA. (A) Influence of ISG20 overexpression on the abundance of viral input DNA. HFF/Control, HFF/ISG20 and HFF/ISG20mut were seeded in triplicates, stimulated with doxycycline (500 ng/ml) for 24 h and infected with HCMV (strain AD169, MOI of 0.5). At 6 und 24 hpi, total DNA was extracted from cells and subjected an HCMV IE1-specific qPCR to determine the number of viral genomes. Values were normalized to levels of cellular albumin and are shown as relative mean values derived from three independent experiments +/- SD. Statistical analysis was performed using a one-sample t-test. Ns, not significant. (B) Influence of ISG20 knockdown on the abundance of viral input DNA. HFF cells were transfected with an ISG20-specific siRNA or a control siRNA and, 72 hours post-transfection, were infected with HCMV (strain AD169, MOI of 0.5). At 6 und 24 hpi, intracellular HCMV DNA was quantified as described in (A). Statistical analysis was performed using a one-sample t-test. Ns, not significant. (C) Influence of ISG20 overexpression on the abundance of newly synthesized viral DNA. HFF/Control, HFF/ISG20 and HFF/ISG20mut were seeded in triplicates, stimulated with doxycycline (500 ng/ml) for 24 h, before they were infected with HCMV (strain AD169, MOI of 0.5). At 72 hpi, intracellular HCMV DNA was quantified as described in (A). Statistical analysis was performed using a one-sample t-test. *, p<0.5; ns, not significant. (D) Influence of ISG20 knockdown on the abundance of newly synthesized viral DNA. HFF cells were transfected with control siRNA or ISG20-specific siRNA using a final concentration of 20 nM. 2 days after transfection, cells were infected with HCMV (strain AD169, MOI of 0.5). At 72 hpi, intracellular viral DNA was quantified as described in (A). Statistical analysis was performed using a one-sample t-test. *, p<0.5; **, p<0.01. (E) Influence of ISG20 overexpression on the stability of newly synthesized viral DNA. HFF/Control, HFF/ISG20 and HFF/ISG20mut were seeded in triplicates, stimulated with doxycycline (500 ng/ml) for 24 h, before they were infected with HCMV (strain AD169, MOI of 0.5). At 72 hpi, cells were treated with foscarnet at a concentration of 400 µM. At different times, after treatment, intracellular viral DNA was quantified as described in (A). Statistically significant differences were calculated for HFF/ISG20 and HFF/ISG20mut in comparison to HFF/Control using a unpaired two-sample t-test. Ns, not significant. Hpi, hours post-infection.

In a next step, we explored whether ISG20 affects newly synthesized genomes, which need to be linearized for packaging and therefore might be more vulnerable to exonuclease-mediated degradation. For this purpose, HFFs expressing ISG20 or ISG20mut and control HFFs were infected with HCMV (strain AD169, MOI of 0.5) for 72 h to ensure the synthesis of new HCMV genomes. The cells were then treated with viral DNA polymerase inhibitor foscarnet to block further DNA synthesis, and the stability of viral DNA was assessed by measuring intracellular genomes using qPCR at different times after treatment. Quantification of HCMV DNA at the time of foscarnet treatment (72 hpi) revealed an overall reduced amount of HCMV genomes in HFF/ISG20 compared to HFF/ISG20mut and HFF/Control (Figure 5C). Vice versa, siRNA-mediated depletion of ISG20 resulted in increased HCMV genomes numbers at 72 hpi (Figure 5D). However, a similar decay rate was observed in ISG20-expressing and control cells suggesting that ISG20 does not affect the stability of viral DNA (Figure 5E).

Collectively, these results provide evidence that ISG20 does not act by degrading HCMV genomes but rather causes a reduction in viral DNA synthesis.

### ISG20 does not induce the degradation of HCMV RNA

To further characterize the mode of action of ISG20, we investigated a possible degradation of viral RNA, as this has been previously demonstrated for HBV [17]. HFF/ISG20wt, HFF/ISG20mut and control cells were infected with HCMV (strain AD169, MOI of 1) and harvested for RNA isolation at 48 hpi, since the antiviral activity of ISG20 was observed from early times after infection. Analysis of viral mRNA levels by reverse-transcription quantitative PCR (RT-qPCR) revealed significantly reduced mRNA levels of early genes UL35 and UL44 in ISG20-overexpressing cells compared to control cells (Figure 6A). This effect was also visible, albeit to a lesser extent, for late genes UL75, UL76, UL82, and UL99. Expression of ISG20mut also led to a reduction in mRNA levels of early and late genes, but not to the same degree as the wildtype ISG20 protein (Figure 6A). In contrast to the reduction of early and late RNAs, elevated mRNA levels of immediate-early genes IE1 and US3 were found in HFF/ISG20, while no significant differences were observed for IE2. To examine whether the reduction of early and late genes is due to a direct degradation of viral RNA by the exonuclease ISG20, we determined the half-life of viral transcripts. Quantification of viral mRNA levels at different times after transcription inhibition by actinomycin D did not reveal significant differences in the decay rate of early (UL44) and late (UL75) RNAs (Figure 6B, C). We conclude that ISG20 leads to a reduction of early and late transcripts, to varying extents, but not via a direct degradation of viral RNAs.

**Figure 6.**
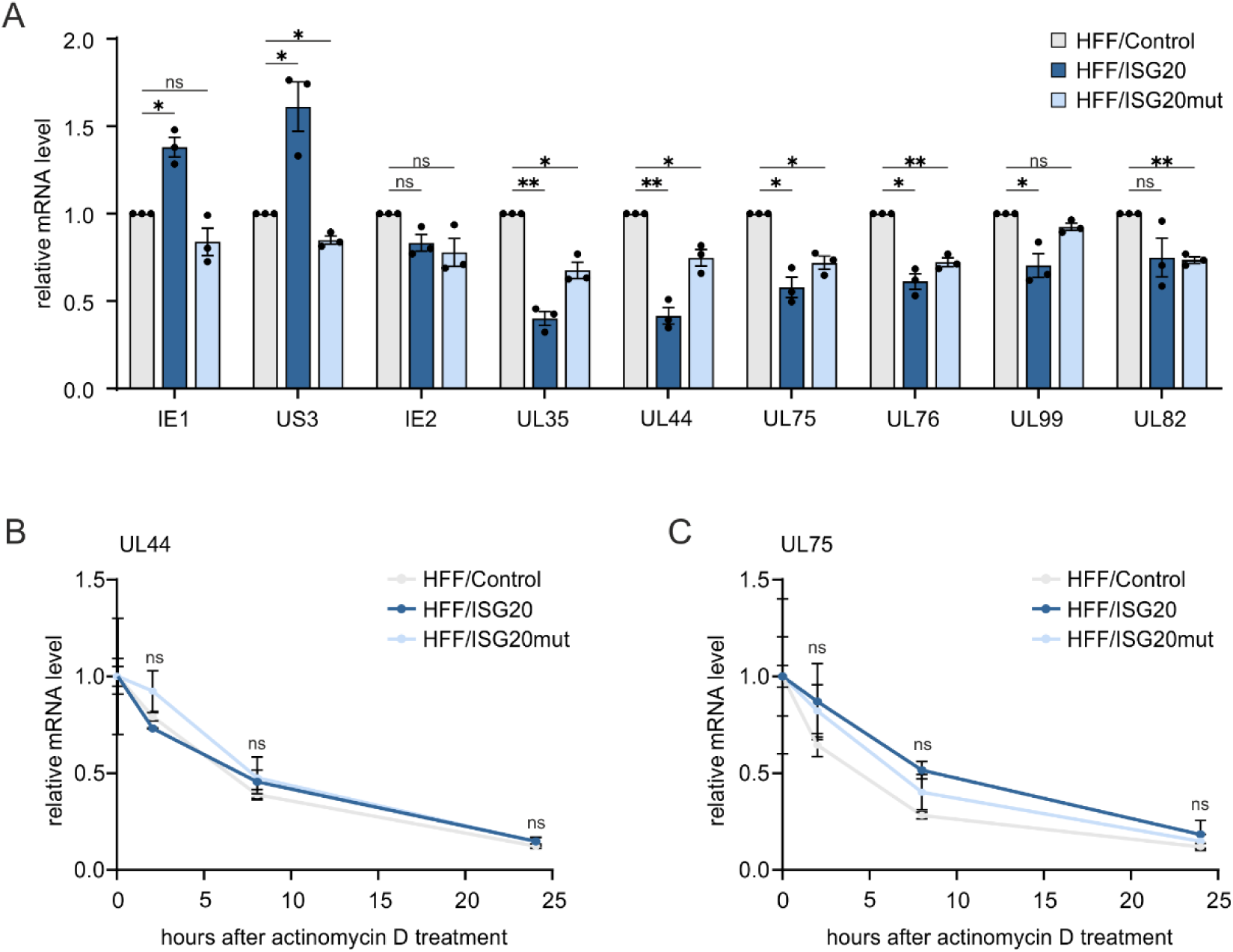
Effect of ISG20 on the stability of HCMV RNA. (A) Influence of ISG20 overexpression on the abundance of viral RNA. HFF/Control, HFF/ISG20 and HFF/ISG20mut were seeded in triplicates, stimulated with doxycycline (500 ng/ml) for 24 h, before they were infected with HCMV (strain AD169, MOI of 1). At 48 hpi, total RNA was extracted from cells and subjected to RT-qPCR to determine viral mRNA levels. Values were normalized to the reference gene GAPDH and are shown as relative mean +/- SD derived from three independent experiments. Statistical analysis was performed using a one-sample t-test. *, p<0.05; **, p<0.01; ns, not significant. (B, C) Influence of ISG20 overexpression on viral RNA decay. HFF/Control, HFF/ISG20 and HFF/ISG20mut were seeded in triplicates, stimulated with doxycycline (500 ng/ml) for 24 h, before they were infected with HCMV (strain AD169, MOI of 1). At 48 hpi, cells were treated with actinomycin D at a final concentration of 2 µg/ml, to block further RNA synthesis. 0 h, 2 h, 8 h, and 24 h after treatment, total RNA was extracted from cells and subjected to RT-qPCR to determine mRNA levels of viral UL44 (B) and UL75 (C). Values were normalized to the reference gene GAPDH and are shown as relative mean values +/- SD calculated from triplicate samples. One representative experiment out of two is shown. Statistical analysis was performed using a unpaired two-tailed t-tests. Ns, not significant.

### ISG20 enhances the induction of interferon-stimulated genes (ISGs)

Besides degrading viral RNA and DNA, ISG20 has been proposed to positively regulate interferon (IFN) signaling, leading to increased expression of IFN-stimulated genes (ISGs) such as IFIT1 and thereby to translation inhibition [19, 20]. In order to examine whether ISG20 enhances ISG induction during HCMV infection, HFF/control, HFF/ISG20wt and HFF/ISG20mut were infected (HCMV strain AD169, MOI of 1) and harvested at 6 hpi, before ISG expression is blocked by IE1 and other viral proteins. Indeed, RT-qPCR analysis revealed an increased expression of IFN-β and all tested ISGs in HFF/ISG20, but not of the upstream transcription factor *IRF3*, which was included as control (Figure 7A). HFFs expressing kinase-dead ISG20mut displayed ISG transcript levels comparable to that of control HFFs. The positive effect of ISG20 on IFN signaling was not only observed in HCMV-infected cells but also upon stimulation of the cGAS-STING DNA sensing pathway with 2’3’-cGAMP (Figure 7B) and upon treatment with IFN-β (Figure 7C). Vice versa, analysis of ISG transcripts after IFN-β treatment of cells with siRNA-mediated depletion of ISG20 revealed a weakened ISG upregulation in absence of ISG20, thereby excluding a non-specific effect of doxycycline-induced ISG20 overexpression (Figure 7D). We further observed that ISG20 expression leads to an increase of ISG transcript levels in the absence of any other stimulus (Figure 7E). Increased ISG abundance in HFF/ISG20 compared to control cells was also observed on protein level, as shown by Western Blot analysis of IFIT1 and IFIT2 expression (Figure 7F). Taken together, our data point towards an antiviral activity of ISG20 that is based on the upregulation of other antiviral ISGs.

**Figure 7.**
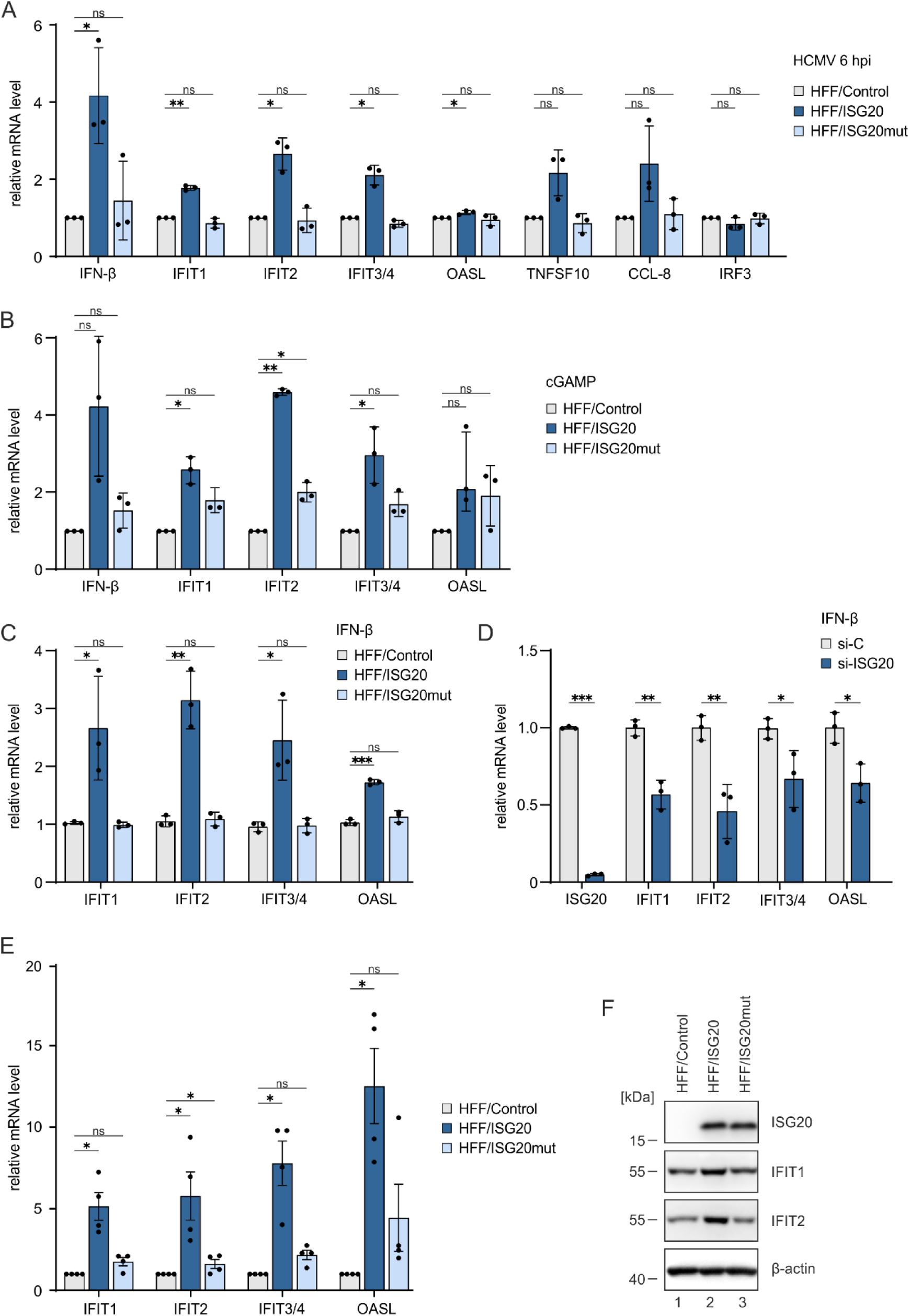
Effect of ISG20 on the expression of IFN and ISGs. (A, B) Influence of ISG20 overexpression on ISG induction upon HCMV infection or cGAMP stimulation. HFF/Control, HFF/ISG20 and HFF/ISG20mut were stimulated with doxycycline (500 ng/ml) for 24 h, before they were infected with HCMV (strain AD169, MOI of 1) for 6h (A) or stimulated with 50 µg/ml 2’3’-cGAMP for 2 h (B). Levels of IFN-β and ISGs were determined by RT-qPCR. Values were normalized to the reference gene GAPDH and are shown as relative mean values +/- SD derived from three independent experiments performed in triplicates. Statistical analysis was performed using a one-sample t-test. *, p<0.05; **, p<0.01; ns, not significant. (C) Influence of ISG20 overexpression on ISG induction by IFN-β. HFF/Control, HFF/ISG20 and HFF/ISG20mut were stimulated with doxycycline (500 ng/ml) for 24 h, before they were treated with 1000 U/ml IFN-β for 6 h. ISG mRNA levels were determined by RT-qPCR. Values were normalized to the reference gene GAPDH and are shown as relative mean values +/- SD derived from triplicate samples. One of two representative experiments is shown. Statistical analysis was performed using an unpaired, two-tailed t-tests. *, p<0.05; **, p<0.01; ***, p<0.001; ns, not significant. (D) Influence of ISG20 knockdown on ISG induction by IFN-β. HFF were transfected with control siRNA (si-C) or ISG20-specific siRNA (si-ISG20) and, after two days, stimulated with 1000 U/ml IFN-β for 6 h. ISG mRNA levels were determined by RT-qPCR. Values were normalized to the reference gene GAPDH and are shown as relative mean values +/- SD derived from triplicate samples. One of two representative experiments is shown. Statistical analysis was performed using an unpaired, two-tailed t-test. *, p<0.05; **, p<0.01; ***, p<0.001. (E) Influence of ISG20 overexpression on ISG induction. HFF/Control, HFF/ISG20 and HFF/ISG20mut were stimulated with doxycycline (500 ng/ml) for 24 h, before they were harvested and subjected to RT-qPCR. Values were normalized to the reference gene GAPDH and are shown as relative mean values +/- SD derived from four independent experiments performed in triplicates. Statistical analysis was performed using a one-sample t-test. *, p<0.05; ns, not significant. (F) Influence of ISG20 overexpression on ISG induction. HFF/Control, HFF/ISG20 and HFF/ISG20mut were stimulated with doxycycline (500 ng/ml) for 48 h, before they were harvested for western blot analysis of ISG20, IFIT1 and IFIT2 expression. Β-actin was included as loading control.

### ISG20 overexpression induces an antiviral gene signature in primary human fibroblasts

To characterize the transcriptomic response of HFF to ISG20 overexpression in more detail, we performed RNA-sequencing (RNA-seq) on RNAs extracted from HFF/ISG20 and HFF/Control. We identified about 1500 differentially expressed genes (DEGs), of which 700 were upregulated and 844 were downregulated in HFF/ISG20. Several ISGs were detected among the upregulated genes, in line with the above results from RT-qPCR analysis (Figure 8A). In addition, we detected an upregulation of IRF9 and XAF1, which are both known for their broad effects on innate immune signaling (Figure 8A). While IRF9, as part of ISGF3, regulates downstream expression of ISGs but also functions beyond the canonical JAK-STAT pathway, XAF1 has recently been described to promote antiviral immune responses by regulating chromatin accessibility [27, 28]. Since epigenetic remodeling by XAF1 acts via degradation of TRIM28 and loss of TRIM28 is known to derepress transposable elements (TE), we asked the question whether ISG20 might also affect the expression of TEs [28, 29]. Indeed, re-analysis of the RNA-seq dataset derived from HFF/ISG20 and HFF/Control cells revealed a significantly increased expression of numerous TEs of all subclasses in ISG20 positive cells (Figure 8B). Functional enrichment analysis of DEGs further revealed that, besides genes associated with IFN and cytokine signaling, genes involved in gene expression and transcription are upregulated upon ISG20 overexpression (Figure 8C). Notably, 10% of all upregulated genes coded for zinc-finger proteins (ZNFss) comprising mainly KRAB domain-containing ZNFss, a group of strong transcriptional repressors involved in cancer-related processes like cell proliferation, differentiation and apoptosis [30] (Figure 8D). However, recent studies suggest that these proteins are also important for antiviral defense [31, 32]. In accordance with a higher expression of KRAB-ZNFs in HFF/ISG20, a downregulation of extracellular matrix-related genes was observed, which recently have been suggested by Ito and colleagues to be suppressed by KRAB-ZNFs in various tumors leading to alterations of cancer phenotypes (Figure 8C and D) [33]. Interestingly, Ito et al. also described that epigenetic activation of human endogenous retroviruses (HERVs) leads to the induction of KRAB-ZNFss, suggestion an interrelation between the upregulation of TEs and KRAB-ZNFs in ISG20 positive cells (Figure 8B and D). Analysis of two existing RNA-seq datasets originating from murine fibroblasts and HEK293T cells expressing ISG20 also showed a high proportion of KRAB-ZNFs among the upregulated genes, indicating a cell type- and species-independent effect of ISG20 [19, 34] (Supplementary Figure 4). Interestingly, analysis of the promoter regions of the upregulated KRAB-ZNFs revealed an enrichment of a binding motif for transcription factors of the ETS-family consisting of a GGAA/T core consensus and additional flanking nucleotides, suggesting a common mechanism of regulation [35] (Figure 8E).

**Figure 8.**
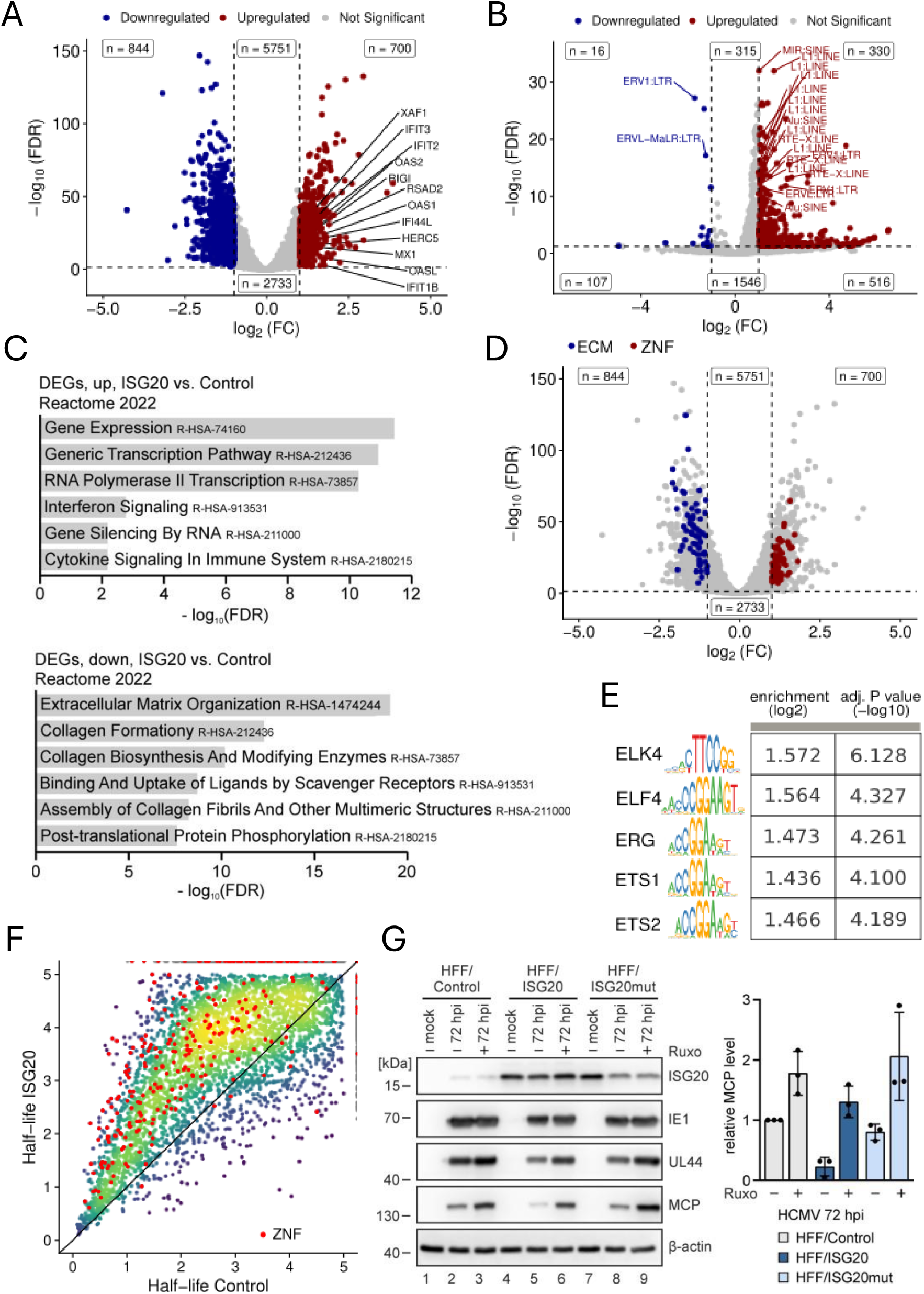
Effect of ISG20 on the cellular transcriptome. (A-D) Transcriptomic changes in ISG20-overexpressing cells analyzed by RNA-seq. HFF/ISG20 and HFF/Control were treated for 24 h with doxycycline (500 ng/ml), before total RNA was isolated and subjected to mRNA sequencing. Results are derived from three biological replicates. (A) Volcano plot showing differentially expressed genes in HFF/ISG20 vs. HFF/Control. Colored dots represent genes with adjusted p-value <0.05 and |log2 fold change) >1. Blue dots represent down-regulated genes, red dots represent up-regulated genes. (B) Volcano plot depicting upregulated TEs in HFF cells expressing ISG20. Red dots represent significantly upregulated TEs from the TEs analysis (log2(FC)>1, P<0.05). (C) Functional enrichment analysis of upregulated genes (upper panel) and downregulated genes (lower panel) using the Reactome 2022 database. (D) Volcano plot showing differentially expressed genes in HFF/ISG20 vs. HFF/Control. Colored dots represent genes with adjusted p-value <0.05 and |log2 fold change) >1. Genes associated with extracellular matrix organization are colored in blue, ZNFs are colored in red. (E) Motif enrichment results from monaLisa, comparing upregulated ZNF promoters against random sequences. Sequence logos represent the enriched motifs, with corresponding log2 enrichment values and -log10 adjusted p-values rounded to three digits in the table. (F) Changes in cellular expression dynamics in HFF/ISG20 analyzed by SLAM-seq. HFF/ISG20 and HFF/Control were treated for 24 h with doxycycline (500 ng/ml) and labeled with 4-thiouridine (400 µM) for 1 h prior to cell lysis, followed by isolation of total RNA. SLAM-seq was performed to identify newly synthesized and total RNA using GRAND-SLAM. RNA-half-lives were estimated using the Bayesian hierarchical model of grandR. Shown are the RNA half-lives in control cells and in ISG20 overexpressing cells. ZNFs are indicated. Axes were deliberately cut at 5 h since longer half-lives cannot be estimated accurately with 1 h RNA labeling. (G) Role of IFN signaling for the antiviral activity of ISG20. HFF/Control, HFF/ISG20 and HFF/ISG20mut were stimulated with doxycycline (500 ng/ml) and treated with 3 µM ruxolitinib (+) or DMSO as control (-), before they were infected with HCMV (strain AD169, MOI of 1) for 72 h and harvested for western blot analysis (left panel). Densitometric analysis of western blots was performed using ImageJ (right panel). Levels of MCP were normalized to β-actin and are shown as relative mean values +/- SD derived from three experiments. ECM, extracellular matrix; DEGs, differentially expressed genes; FC, fold change; FDR, false discovery rate; ZNFs, zinc finger proteins; ruxo, ruxolitinib.

Having observed that ISG20 overexpression stimulates the expression of genes associated with viral infection and tumorigenesis, we next analyzed the effect on the stability of cellular transcripts, as the exonuclease domain has been shown to be important for ISG20 function. For this, newly synthesized RNA in HFF/ISG20 and HFF/Control was labeled for 1 h with 4-thiouridine (4sU) followed by thiol-linked alkylation for the metabolic sequencing of RNA (SLAM-seq) [36]. Newly synthesized and total RNA were then identified using the computational approach GRAND-SLAM [37]. Surprisingly, a general stabilization of cellular transcripts in HFF/ISG20 was observed, as recognized from the overall shift of the new-to-total RNA ratio (Figure 8F). This finding indicates that ISG20 does not act via a destabilization of mRNAs encoding regulatory proteins.

Finally, to substantiate the importance of ISG20-mediated perturbation of the cellular transcriptome for HCMV replication, we sought for possibilities to interfere with cellular gene functions. Since we hypothesize that ISG20 inhibits HCMV replication through inducing antiviral genes such as ISGs, we analyzed HCMV infection in the presence of an inhibitor of IFN signaling, the JAK inhibitor ruxolitinib (Figure 8G). As observed in previous experiments (see Figure 4), we detected a negative impact of ISG20 on the expression of viral early (UL44) and late (MCP) genes at 72 hpi. Treatment with ruxolitinib, however, restored the ability of HCMV to replicate efficiently in ISG20-expressing HFF, as determined by measuring the abundance of the late phase protein MCP (Figure 8G, right panel). In conclusion, our data provide evidence that ISG20 induces an antiviral gene signature in HFF cells, thereby interfering with HCMV replication.

## Discussion

The interferon inducible 20 kDa protein ISG20 is best known for its inhibitory activity on a broad spectrum of unrelated RNA viruses including members of the *Retroviridae, Orthomyxoviridae, Rhabdoviridae, Picornaviridae* and *Flaviviridae* families [10, 11, 38, 39]. For these viruses, degradation of viral RNA is supposed to serve as the main mechanism of antiviral action [3]. Consistent with this, ISG20 is classified as a member of the DEDD subgroup of 3’-5’ exonucleases, which contain three conserved exonuclease motifs, and in vitro experiments have demonstrated a strong degradative activity for single-stranded nucleic acid substrates with a preference for RNA [9, 40]. Indeed, recent studies revealed specific RNA secondary structures in the 3’ untranslated region of the Usutu flavivirus that confer resistance to degradation thus emphasizing the relevance of ISG20’s exonuclease activity for the inhibition of RNA viruses [41].

Here, we demonstrate that ISG20 acts antiviral against herpesviruses. We identified *ISG20* in a single cell RNA sequencing experiment within a gene signature of primary human fibroblasts that correlated with restriction of human cytomegalovirus (HCMV) gene expression (Figure 1A-C). Investigation of the antiviral activity of members of this gene signature utilizing an siRNA screen detected a clear proviral effect after depletion of ISG20 (Figure 1D). Evidence for its contribution to the host cell’s defense against human cytomegalovirus infection is also provided by the observation that ISG20 is transiently upregulated during HCMV infection (Figure 2B) and counteracted by the interferon-antagonistic immediate-early protein IE1 (Figure 2C). Furthermore, a clear delay in the replication of various strains of human cytomegalovirus and of herpes simplex virus type I was detected by multistep growth curve analyses after infection of primary human fibroblasts with doxycycline-inducible expression of ISG20 (Figure 3). This indicates that both alpha- and beta-herpesviruses are amenable to inhibition by ISG20. Interestingly, for Kaposi-sarcoma associated herpesvirus (KSHV) ISG20 was proposed to play a role for the maintenance of viral latency, suggesting that ISG20 modulates the gene expression program of all subclasses of the herpesviruses [42].

Overexpression as well as knockdown of ISG20 primarily affected the accumulation of early and late viral proteins indicating that the inhibition of HCMV gene expression occurs at a step beyond the immediate early phase of the replication cycle (Figure 4). This argues against the degradation of incoming circularized HCMV DNA by ISG20. This appeared as a potential antiviral mechanism since a recent publication described the degradation of covalently closed circular DNA by ISG20 which was observed in the context of persistent hepatitis B virus infection [18]. Consistently, we neither detected a negative impact of ISG20 on the amount of HCMV input genomes nor on newly synthesized genomes thus excluding a degradation of HCMV DNA by ISG20 (Figure 5).

Due to the known propensity of ISG20 to degrade RNA, we speculated that early and late HCMV transcripts, which are predominantly intronless and require a viral RNA export factor for transport to the cytoplasm, may serve as the target [43]. For that reason, we carefully measured the half-lives of prototypical early and late viral transcripts in the absence or presence of ISG20, however, were not able to observe a significant difference in the decay rate (Figure 6B and C). Moreover, RNA half-lives of cellular transcripts were determined by SLAM-seq and subsequent GRAND-SLAM analysis [37] (Figure 8E). Surprisingly, ISG20 expression elicited a general stabilization of cellular transcripts, arguing against a broad degradative effect of ISG20 on RNA. However, we cannot exclude that ISG20 affects the expression of microRNAs or long noncoding RNAs, which could influence the expression of many different genes [44]. For instance, ISG20’s effect on KSHV latency was proposed to be due to the regulation of viral microRNAs [42].

Correlating with the stabilization of cellular mRNAs our transcriptomic analysis revealed the upregulation of three distinct gene sets that altogether indicate the induction of a so far undescribed innate immune defense signature by ISG20 in primary human fibroblast. Firstly, a distinct set of cytokines, ISGs and ISG-regulatory factors with well-known functions in antiviral defense was upregulated (Figure 8A). This included the oligoadenylate synthase (OAS) 1, the myxovirus resistance protein MX1 and the interferon induced protein with tetratricopeptide repeats IFIT1, for which strong anti-HCMV effects have been reported [45]. In addition, two factors known for their expanded capacity on innate immune signaling were found to be elevated: (i) IRF9, as part of ISGF3, is important for ISG activation upon JAK-STAT signaling but also exerts functions independent of the canonical pathway and (ii) XAF1, which is supposed to promote antiviral responses by regulating chromatin accessibility [27, 28].

Of note, upregulation of type I interferon response proteins by ISG20 has previously been described in the context of the RNA alphaviruses Chikungunya (CHIKV) and Venezuelan equine encephalitis virus (VEEV) [19]. Ectopic expression of ISG20 in murine embryonic fibroblasts strongly inhibited the replication of both CHIKV and VEEV without any evidence for RNA degradation which is consistent with our results. Instead, a suite of >100 genes with predicted antiviral activity, including many ISGs, was induced by ISG20 [19]. Replication inhibition was due to a block of alphavirus genome translation which resulted from strong upregulation of IFIT1 by ISG20 [19]. Although IFIT1 might also contribute to ISG20’s interference with HCMV replication [45], translation inhibition appears unlikely since the downregulation of viral early/late RNAs and proteins correlated well (Figures 4 and 6A).

Consistent with our study, Weiss et al. also observed that a mutation of ISG20 within the Exo II domain, which disrupts the exonuclease activity, alleviates the antiviral effects [19] (Figure 3). Although this suggests a requirement of ISG20’s enzymatic activity for antiviral function, the respective mutation in exo II might also affect other protein interactions of ISG20 that are required for activity.

A second suite of upregulated genes detected by RNA-seq analysis of ISG20-expressing HFFs encoded zinc-finger proteins (ZNFs), in particular Krüppel associated box (KRAB) domain containing ZNFs (Figure 8D). KRAB-ZNFs are best known for their propensity to act as strong epigenetic repressors, however, the functions of most members of this large protein family remain unknown [46]. A reanalysis of existing data sets for ISG20-expressing murine embryonic fibroblasts and 293T cells detected a similar upregulation of KRAB-ZNFs, indicating that ISG20 elicits this expression signature in a cell-type independent manner (Supplementary Figure 4). Of note, enhanced expression of KRAB-ZNFs and ISG20 occurs in a number of different tumors [20, 47, 48]. Ito et al. recently described a transcriptome signature of tumor cells, that comprises a co-regulation of human endogenous retroviruses (HERVs) and KRAB-ZNFs, which were upregulated, and extracellular matrix (ECM) genes, which were downregulated [33]. We could also detect a broad induction of transposable elements (TEs) including HERVs as well as a downregulation of ECMs in ISG20 expressing primary human fibroblasts (Figure 8 B and D). This suggests a common gene-regulatory network that co-regulates TEs, KRAB-ZNFs and ECMs. While ECMs seem to be repressed by KRAB-ZNFs, the co-induction of TEs and KRAB-ZNFs may be explained by epigenetic derepression of TEs which, in turn, stimulate the expression of KRAB-ZNF genes that are preferentially encoded nearby HERVs [33]. Interestingly, this gene expression signature correlated with suppressive effects on tumor progression [33]. This was also reported for ISG20 positive ovarian cancer cells and may involve the enhancement of IFN-beta production and response which is known to driven by the upregulation of HERVs [20, 49]

Consistent with the notion, that the ISG20-induced gene signature consisting of upregulated ISGs, ZNFs and TEs augments the IFN-mediated host cell defense we observed that ISG20 enhanced both IFN-beta and ISG expression after infection with HCMV (Figure 7A). This was also detected after either stimulation of the cGAS-STING pathway with cGAMP or after treatment with IFN-beta (Figure 7B and C). Importantly, siRNA mediated depletion of ISG20 led to a considerable decrease of the ISG response after IFN-beta stimulation (Figure 7D). This finally demonstrates that ISG20 serves as an amplifier of the IFN response. Consistently, we detected that treatment with the JAK-STAT inhibitor ruxolitinib alleviated the inhibitory effect of ISG20 on HCMV infection (Figure 8E). Taken together, our data show that ISG20 induces an innate immune defense signature comprising a upregulation of ISGs, KRAB-ZNFs and TEs that is able to amplify the IFN-mediated host cells defense thus explaining its broad antiviral activity.

## Material and methods

### Expression plasmids

Oligonucleotide primers for generation of expression plasmids were purchased from Biomers GmbH (Ulm, Germany) and are listed in Table 2. A pCMV6-Entry plasmid containing the coding sequence for ISG20 (NM_002201) was obtained from Origene, the coding sequence for ISG20 mutant D94G (ISG20mut) from Biocat GmbH gene synthesis service. To generate plasmids for doxycycline-inducible ISG20/ISG20mut expression, both sequences were amplified via PCR with primers listed in Table 2 and inserted into pInducer20-CRSmut [50] by a combined BP/LR Gateway recombination reaction using pDONR221 (Invitrogen) as entry vector. Lentiviral vectors for stable expression of mGFP-tagged ISG20 and ISG20mut were generated by PCR amplification of the ISG20 and ISG20mut sequences using primers specified in Table 2 and insertion into pLenti-EF1a-C-mGFP-P2A-Puro (Origene) using the enzymes SgsI and MluI.

**Table 1.**
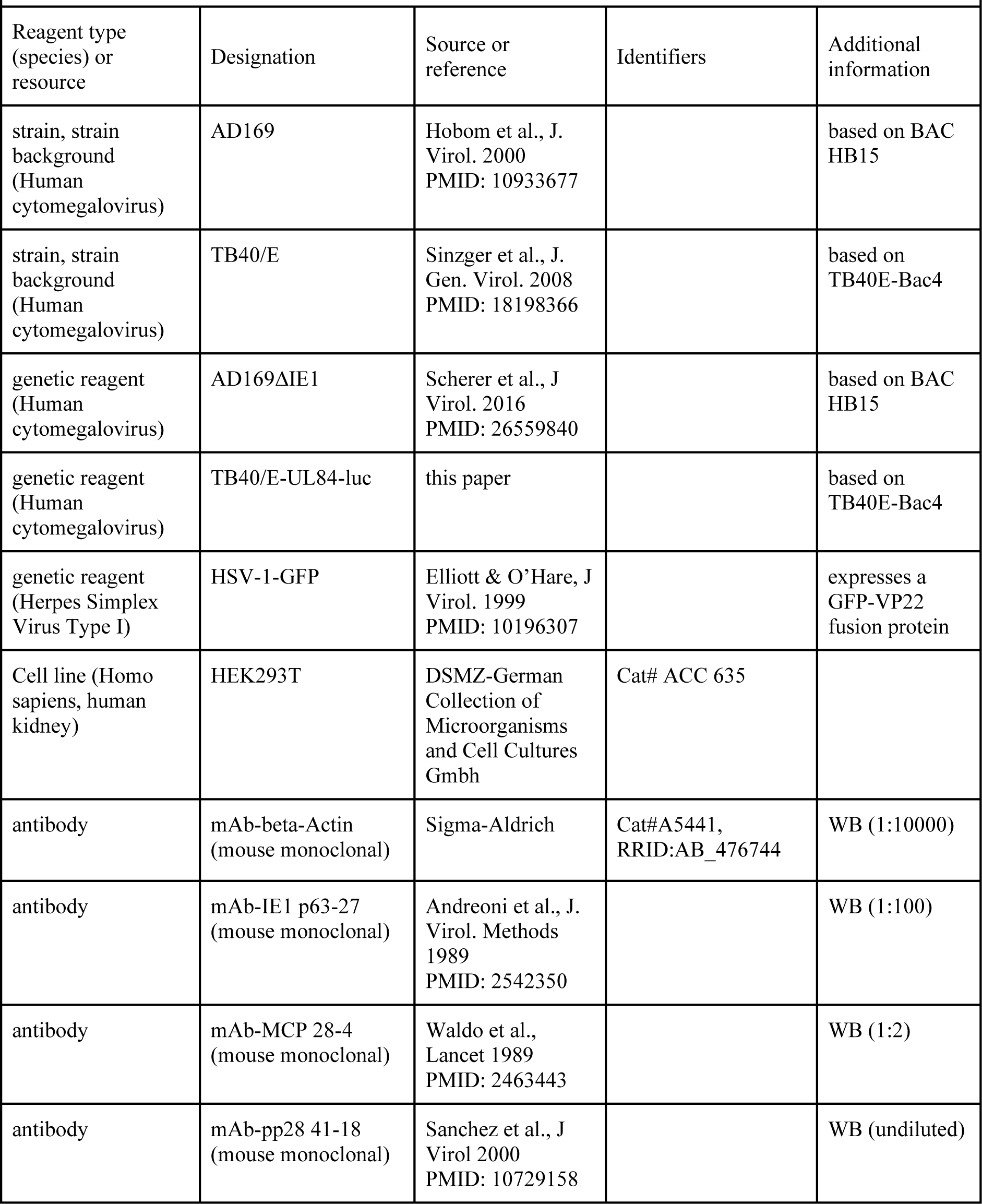
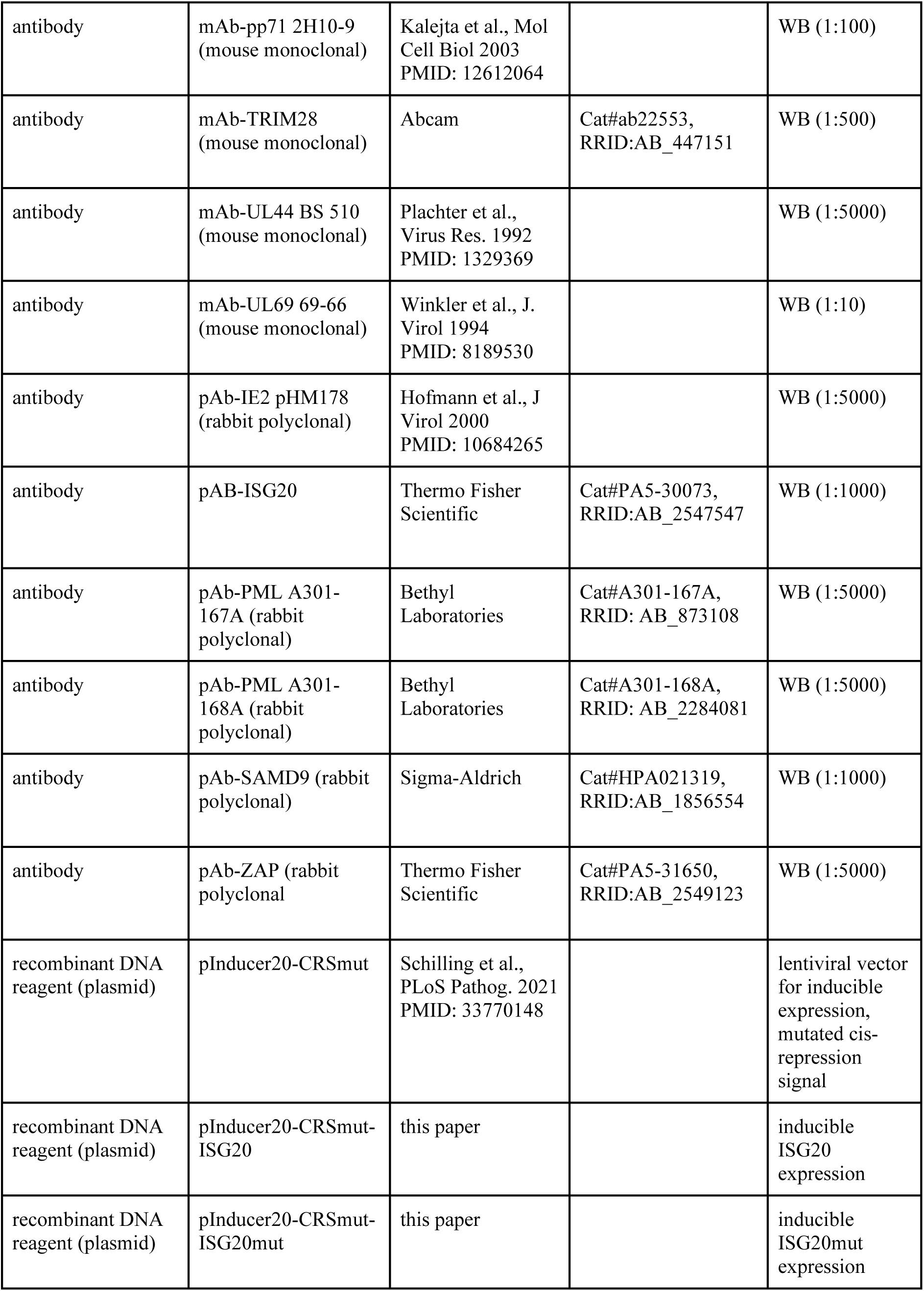
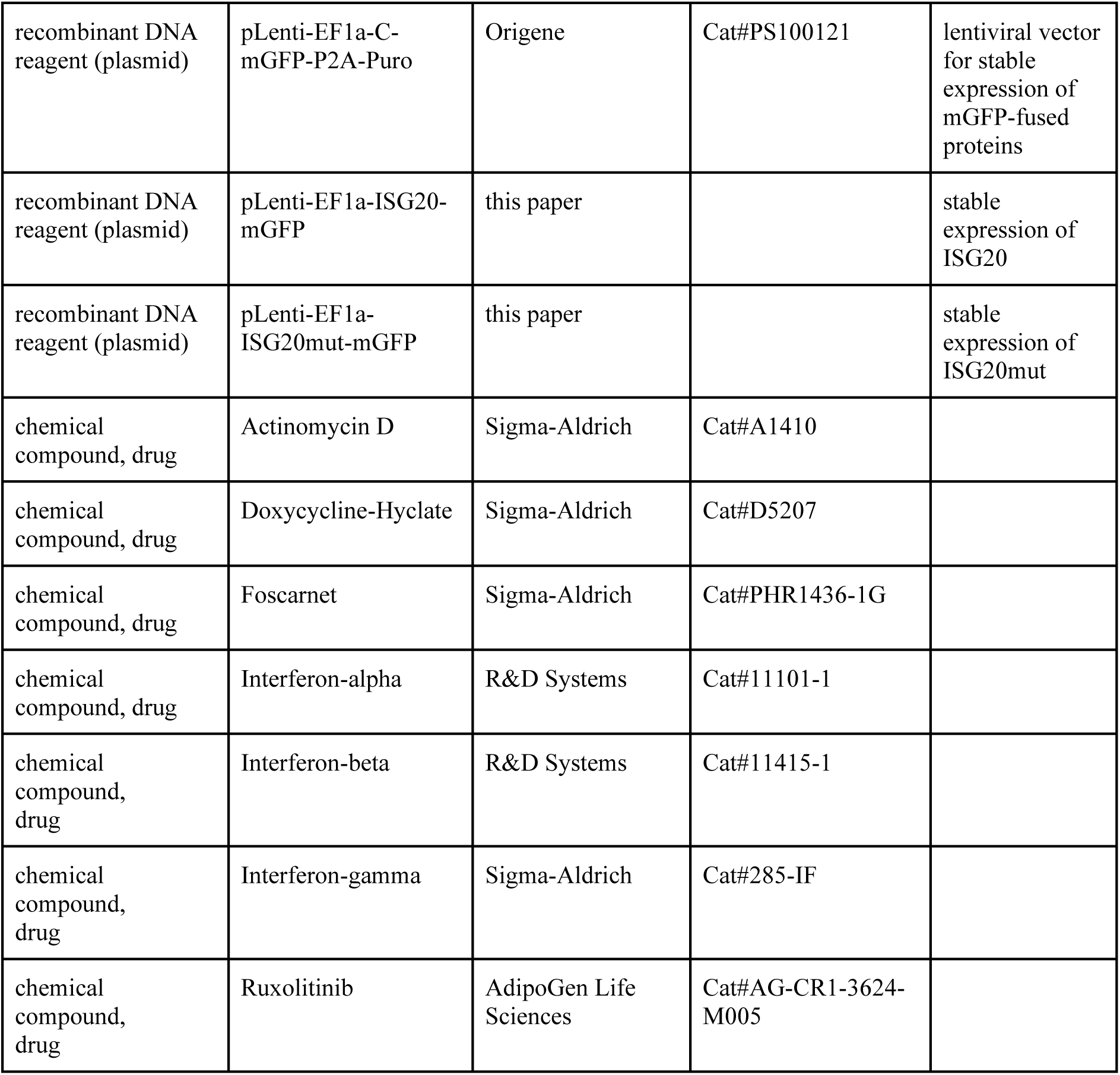
Key resources table.

| Table 1. Key Resources Table |  |  |  |  |
| --- | --- | --- | --- | --- |
| Reagent type (species) or resource | Designation | Source or reference | Identifiers | Additional information |
| strain, strain background (Human cytomegalovirus) | AD169 | Hobom et al., J. Virol. 2000<br>PMID: 10933677 |  | based on BAC HB15 |
| strain, strain background (Human cytomegalovirus) | TB40/E | Sinzger et al., J. Gen. Virol. 2008<br>PMID: 18198366 |  | based on TB40E-Bac4 |
| genetic reagent (Human cytomegalovirus) | AD169ΔIE1 | Scherer et al., J Virol. 2016<br>PMID: 26559840 |  | based on BAC HB15 |
| genetic reagent (Human cytomegalovirus) | TB40/E-UL84-luc | this paper |  | based on TB40E-Bac4 |
| genetic reagent (Herpes Simplex Virus Type I) | HSV-1-GFP | Elliott & O'Hare, J Virol. 1999<br>PMID: 10196307 |  | expresses a GFP-VP22 fusion protein |
| Cell line (Homo sapiens, human kidney) | HEK293T | DSMZ-German Collection of Microorganisms and Cell Cultures Gmbh | Cat# ACC 635 |  |
| antibody | mAb-beta-Actin (mouse monoclonal) | Sigma-Aldrich | Cat#A5441, RRID:AB_476744 | WB (1:10000) |
| antibody | mAb-IE1 p63-27 (mouse monoclonal) | Andreoni et al., J. Virol. Methods 1989<br>PMID: 2542350 |  | WB (1:100) |
| antibody | mAb-MCP 28-4 (mouse monoclonal) | Waldo et al., Lancet 1989<br>PMID: 2463443 |  | WB (1:2) |
| antibody | mAb-pp28 41-18 (mouse monoclonal) | Sanchez et al., J Virol 2000<br>PMID: 10729158 |  | WB (undiluted) |
| antibody | mAb-pp71 2H10-9<br>(mouse monoclonal) | Kalejta et al., Mol<br>Cell Biol 2003<br>PMID: 12612064 |  | WB (1:100) |
| antibody | mAb-TRIM28<br>(mouse monoclonal) | Abcam | Cat#ab22553,<br>RRID:AB_447151 | WB (1:500) |
| antibody | mAb-UL44 BS 510<br>(mouse monoclonal) | Plachter et al.,<br>Virus Res. 1992<br>PMID: 1329369 |  | WB (1:5000) |
| antibody | mAb-UL69 69-66<br>(mouse monoclonal) | Winkler et al., J.<br>Virol 1994<br>PMID: 8189530 |  | WB (1:10) |
| antibody | pAb-IE2 pHM178<br>(rabbit polyclonal) | Hofmann et al., J<br>Virol 2000<br>PMID: 10684265 |  | WB (1:5000) |
| antibody | pAB-ISG20 | Thermo Fisher<br>Scientific | Cat#PA5-30073,<br>RRID:AB_2547547 | WB (1:1000) |
| antibody | pAb-PML A301-167A (rabbit<br>polyclonal) | Bethyl<br>Laboratories | Cat#A301-167A,<br>RRID: AB_873108 | WB (1:5000) |
| antibody | pAb-PML A301-168A (rabbit<br>polyclonal) | Bethyl<br>Laboratories | Cat#A301-168A,<br>RRID: AB_2284081 | WB (1:5000) |
| antibody | pAb-SAMD9 (rabbit<br>polyclonal) | Sigma-Aldrich | Cat#HPA021319,<br>RRID:AB_1856554 | WB (1:1000) |
| antibody | pAb-ZAP (rabbit<br>polyclonal) | Thermo Fisher<br>Scientific | Cat#PA5-31650,<br>RRID:AB_2549123 | WB (1:5000) |
| recombinant DNA<br>reagent (plasmid) | pInducer20-CRSmut | Schilling et al.,<br>PLoS Pathog. 2021<br>PMID: 33770148 |  | lentiviral vector<br>for inducible<br>expression,<br>mutated cis-<br>repression<br>signal |
| recombinant DNA<br>reagent (plasmid) | pInducer20-CRSmut-ISG20 | this paper |  | inducible<br>ISG20<br>expression |
| recombinant DNA<br>reagent (plasmid) | pInducer20-CRSmut-ISG20mut | this paper |  | inducible<br>ISG20mut<br>expression |
| recombinant DNA reagent (plasmid) | pLenti-EF1a-C-mGFP-P2A-Puro | Origene | Cat#PS100121 | lentiviral vector for stable expression of mGFP-fused proteins |
| recombinant DNA reagent (plasmid) | pLenti-EF1a-ISG20-mGFP | this paper |  | stable expression of ISG20 |
| recombinant DNA reagent (plasmid) | pLenti-EF1a-ISG20mut-mGFP | this paper |  | stable expression of ISG20mut |
| chemical compound, drug | Actinomycin D | Sigma-Aldrich | Cat#A1410 |  |
| chemical compound, drug | Doxycycline-Hyclate | Sigma-Aldrich | Cat#D5207 |  |
| chemical compound, drug | Foscarnet | Sigma-Aldrich | Cat#PHR1436-1G |  |
| chemical compound, drug | Interferon-alpha | R&D Systems | Cat#11101-1 |  |
| chemical compound, drug | Interferon-beta | R&D Systems | Cat#11415-1 |  |
| chemical compound, drug | Interferon-gamma | Sigma-Aldrich | Cat#285-IF |  |
| chemical compound, drug | Ruxolitinib | AdipoGen Life Sciences | Cat#AG-CR1-3624-M005 |  |

**Table 2:**
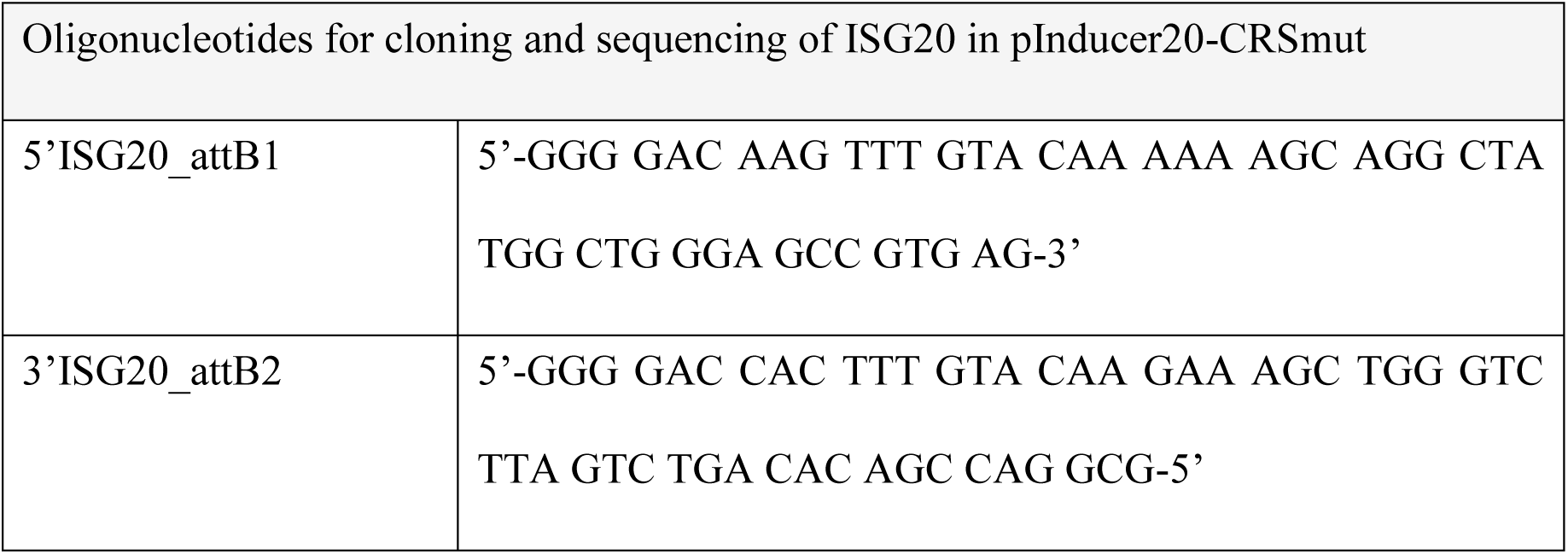
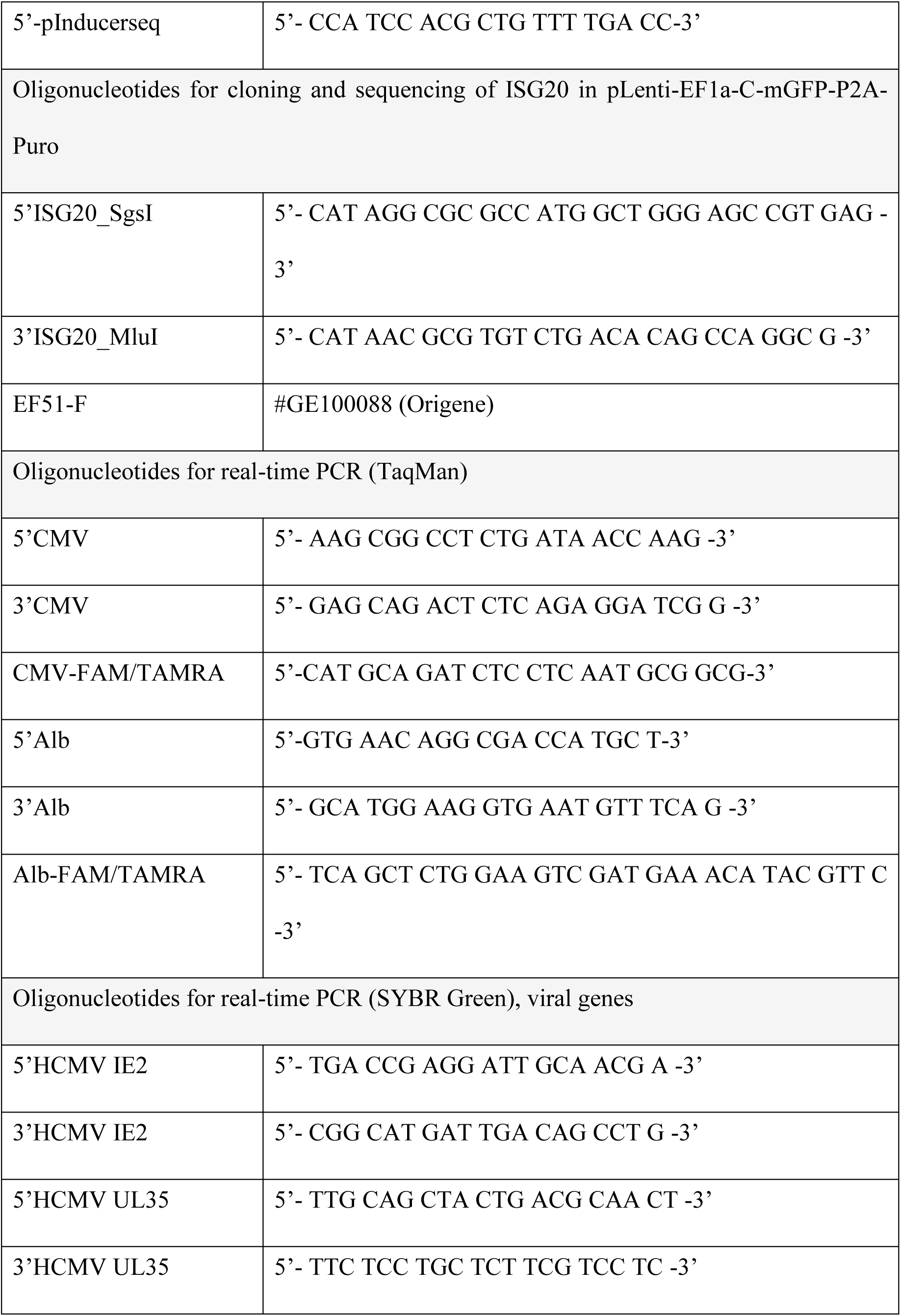
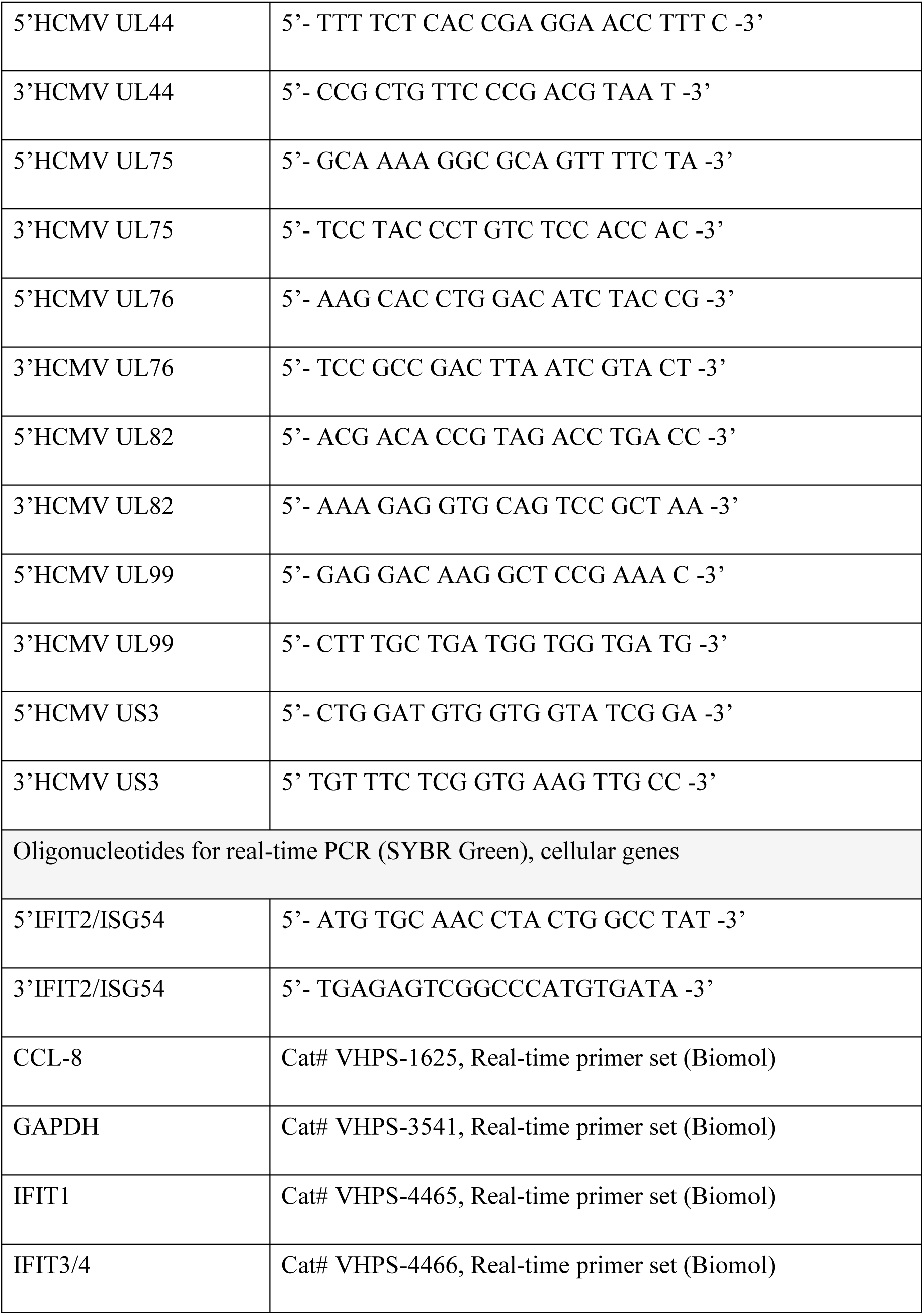
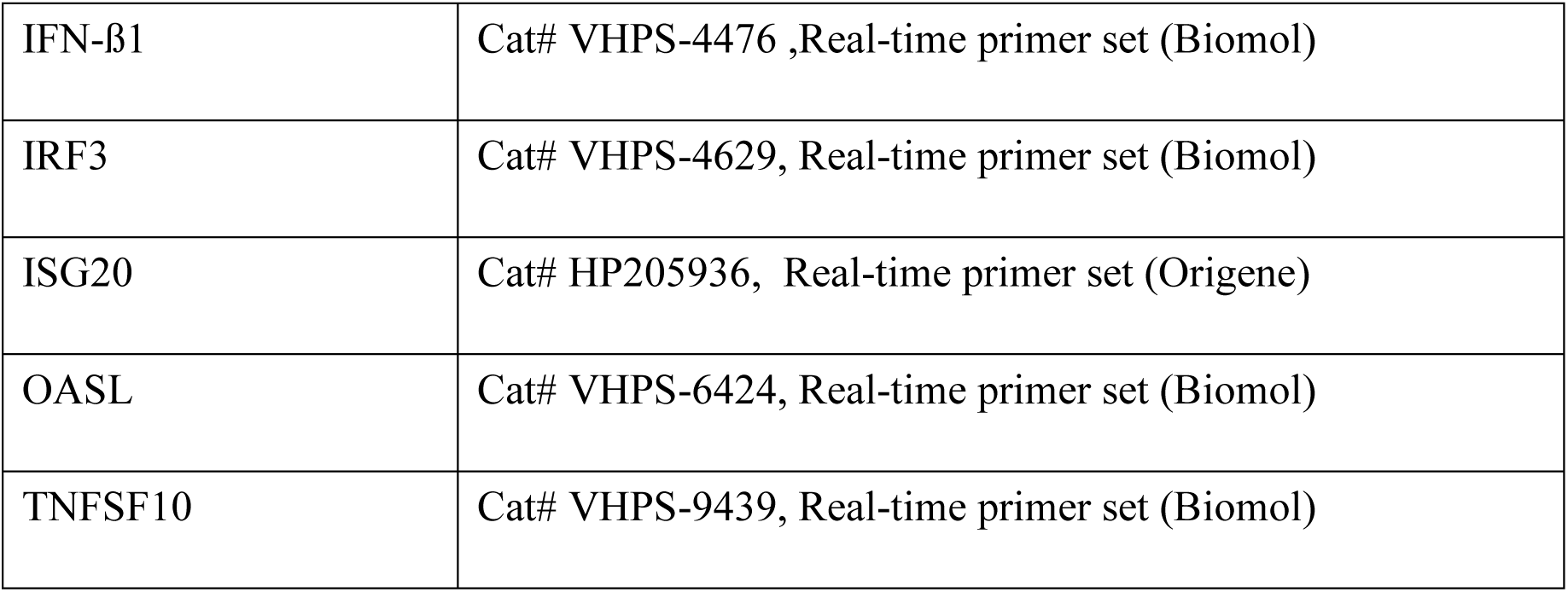
Oligonucleotides.

### Cells and viruses

Primary human foreskin fibroblast (HFF) cells were isolated from human foreskin tissue and cultivated at 37 °C and 5% CO_2_ in Eagle’s minimal essential medium (Gibco) containing 7 % fetal calf serum (FCS) (BIO&SELL), 1% GlutaMAX (Gibco), and penicillin-streptomycin (Sigma-Aldrich). HFF with inducible expression of ISG20 or ISG20mut were cultured in the presence of 500 μg/ml geneticin (InvivoGen) and 10% FCS. HFF with stable expression of ISG20-mGFP or ISG20mut-mGFP were cultured in the presence of 1 µg/ml puromycin (InvivoGen) and 10% FCS. HEK293T (DSMZ ACC 635) cells were cultivated in Dulbecco’s minimal essential medium (DMEM) containing Glutamin (Gibco) and supplemented with 10% FCS (Sigma-Aldrich) and penicillin-streptomycin (Sigma-Aldrich). Cells were tested for absence of mycoplasma contamination using the commercially available MycoAlert mycoplasma detection kit (Lonza).

Infection experiments of HFF cells were performed with the HCMV strains TB40/E and AD169, as well as recombinant viruses TB40/E-UL84-luc and AD169ΔIE1 [51]. Reconstituted HSV-1 expressing a GFP-tagged VP22 was kindly provided by Manfred Marschall (Institute for Clinical and Molecular Virology, Erlangen, Germany) [52].

For titration of HCMV, HFF were infected with serial dilutions of virus supernatants and, after 24 h of incubation, cells were fixed and stained for IE1. IE1-deleted HCMV was titrated by staining of IE2. Automated quantification of IE1/IE2-positive cells was performed using the ImageXpress Pico System (Molecular Devices), and viral titers were calculated in immediate-early (IE) units/ml. Titration of HSV-1-GFP was conducted by infection HFF with virus dilutions and 16 hpi, determination GFP-positive cells by FACS analysis using a CytoFLEX SRT Cell Sorter (Beckman Coulter). The viral titer was calculated in GFP units/ml.

For construction of the TB40/E-UL84-luc BAC by homologous recombination, a linear recombination fragment was isolated from a transfer plasmid, comprising firefly luciferase (derived from the pGFL-Basic vector, Promega) under control of a UL84 promotor as well as a kanamycin cassette marker flanked by the HCMV genes US9 and US10. This fragment was used for electroporation of competent *Escherichia coli* strain DH10B harboring TB40/E BAC4 and the plasmid pKD46. Subsequent recombination was performed as described previously in order to insert UL84-luc between ORFs US9 and US10 [53]. The integrity of the resulting recombinant BAC was confirmed by PCR, restriction enzyme digestion and direct sequencing.

### Lentiviral transduction of HFF cells

Lentiviral transduction was used to establish HFF with inducible or stable expression of ISG20 or ISG20mut (D94G). Replication-deficient lentiviruses were produced in HEK293T cells, which were seeded in 10 cm dishes at a density of 5 × 10^6^ cells/dish. After 24 h, cells were transfected with the respective lentiviral vector together with packaging plasmids pLP1, pLP2, and pLP/VSV-G using the Lipofectamine 2000 reagent (Invitrogen). 16 h later, cells were provided with fresh medium and 48 h after transfection, viral supernatants were harvested, filtered through a 0.45 µm sterile filter, and stored at −80 °C. To transduce primary HFF, 8 x 10^4^ cells/well were seeded in 6-well plates. One day later, the cells were incubated with lentivirus supernatant in the presence of 7.5 µg/ml polybrene (Sigma-Aldrich) for 24 h. Stably transduced cell populations were selected by adding 500 µg/ml geneticin (doxycycline-inducible expression of ISG20) or 1 µg/ml puromycin (stable expression of ISG20-mGFP) to the cell culture medium.

### SiRNA transfection of HFF cells

HFF cells were seeded in 96-well plates at a density of 1 × 10^4^ cells/well or in 6-well plates at a density of 2.8 × 10^5^ cells/well. The next day, the cell culture medium in each well was replaced with 50 µl (96-well format) or 500 µl (6-well format) Opti-MEM (Gibco). Subsequently, 2 pmol (96-well format) or 50 pmol (6-well format) of the respective siRNA were transfected into the HFF cells using Lipofectamine 2000 or Lipofectamine RNAiMax (Invitrogen) according to the manufacturer’s instructions. Infection or IFN-treatment of the cells was conducted at 2 to 3 days post-transfection. Utilized siRNAs are listed in Table 3.

**Table 3.** ON-TARGETplus siRNA Smart Pools (Horizon Discovery).

| Gene | Gene ID | Gene Accession | Cat# |
| --- | --- | --- | --- |
| *APOL2* | 23780 | NM_030882 | L-017407-00 |
| *APOL6* | 80830 | NM_030641 | L-013432-01 |
| *DDX58* | 23586 | NM_014314 | L-012511-00 |
| *DDX60* | 55601 | NM_017631 | L-017664-01 |
| *FGF2* | 2247 | NM_002006 | L-006695-00 |
| *HERC5* | 51191 | NM_016323 | L-005174-00 |
| *HERC6* | 55008 | NM_001013000 | L-005175-00 |
| *HIP1R* | 9026 | NM_003959 | L-027079-00 |
| *IFI44* | 10561 | NM_006417 | L-016368-00 |
| *IFIH1* | 64135 | NM_022168 | L-013041-00 |
| *IFIT1* | 3434 | NM_001001887 | L-019616-00 |
| *IFIT2* | 3433 | NM_001547 | L-012582-02 |
| *IFIT3* | 3437 | NM_001549 | L-017691-00 |
| *ISG15* | 9636 | NM_005101 | L-004235-03 |
| *ISG20* | 3669 | NM_002201 | L-015994-00 |
| *MX1* | 4599 | NM_002462 | L-011735-00 |
| *MX2* | 4600 | NM_002463 | L-011736-00 |
| *OAS2* | 4939 | NM_001032731 | L-009768-00 |
| *OASL* | 8638 | NM_198213 | L-012617-00 |
| *PARP14* | 54625 | NM_017554 | L-023583-00 |
| *PMAIP1* | 5366 | NM_021127 | L-005275-00 |
| *RARRES3* | 5920 | NM_004585 | L-011889-00 |
| *RNF149* | 284996 | NM_173647 | L-007169-00 |
| *RSAD2* | 91543 | NM_080657 | L-015423-00 |
| *SAMD9* | 54809 | NM_017654 | L-031141-00 |
| *SAMD9L* | 219285 | NM_152703 | L-028907-00 |
| *WARS* | 7453 | NM_213645 | L-008322-00 |
| *ZC3HAV1* | 56829 | NM_024625 | L-017449-01 |
| *ZFP36L2* | 678 | NM_006887 | L-013605-01 |
| *ZNFX1* | 57169 | NM_021035 | L-014074-00 |
| Non-targeting control pool |  |  | D-001810-10 |

### Cell viability assay

HFF cells were seeded in 96-well plates and treated the next day with doxycycline or siRNA as described above. On the day of analysis, the medium was replaced with 100 µl pre-warmed fresh medium per well. 100 µl of the CellTiterGlo buffer, contained in the CellTiter-Glo Luminescent Cell Viability Assay Kit (Promega), was added to each well and the resulting luminescence was measured in a Plate Chameleon luminometer (Hidex Deelux Labortechnik GmbH) using the software MikroWin2000 (Labsis) or a SpectraMax iD5 Multi-Mode Microplate Reader (Molecular Devices) with the corresponding SoftMax Pro software.

### Western blotting

Lysates from transfected or infected HFF cells, which were seeded in 6-well plates, were prepared in a sodium dodecyl sulfate-polyacrylamide gel electrophoresis (SDS-PAGE) loading buffer by boiling for 10 min at 95 °C and sonication for 1 min. Proteins were separated on sodium dodecyl sulfate-containing 8 to 15% polyacrylamide gels and transferred to PVDF membranes (Biorad), followed by chemiluminescence detection using a FUSION FX7 imaging system (Vilber).

### Luciferase assay to determine viral replication

HFF cells were seeded in clear-bottom 96-well plates and transfected with siRNAs as described above. Cells were infected with TB40/E-UL84-luc at a MOI 0.01, which expresses firefly luciferase under the control of the UL84 early promoter. 7 days after infection, luciferase expression by the virus was measured using the Luc-Screen Extended-Glow Luciferase Reporter Gene Assay System (Applied Biosystems) according to manufacturer’s instructions. The resulting luminescence was measured with a luminometer (Plate Chameleon, Hidex Deelux Labortechnik GmbH) using the software MikroWin2000 (Labsis).

### Quantification of extra- and intracellular HCMV genomes by real-time PCR

For multistep growth curve analysis, HFF cells were seeded in triplicates into 6-well dishes at a density of 3 × 10^5^ cells/well. Inducible cells were treated with doxycycline for one day, before they were infected with HCMV at a MOI of 0.1 or 0.01. Supernatants from infected cells were harvested at indicated days after inoculation and cells were provided with fresh medium containing doxycycline. The supernatants were subjected to proteinase K treatment, before viral genome copies were quantified by real-time PCR.

To quantify intracellular HCMV genomes, infected HFF cells, which were seeded in triplicates into 6-well dishes, were harvested and cell pellets were collected. Cell pellets were washed one time with 1ml PBS and subsequently resuspended in 200 µl PBS. Total DNA was isolated using the DNeasy Blood & Tissue Kit (Qiagen) according to manufacturer’s instructions. Viral genome copies were quantified by real-time PCR and normalized to cellular albumin copies. Real-time PCR was conducted using an Agilent AriaMx Real-time PCR System and the analysis software Agilent Aria 1.5 (Agilent Technologies, Inc.). HCMV genomes were quantified by amplification of an HCMV IE1-(UL123-) specific target sequence with primers 5’CMV and 3’CMV along with the hydrolysis probe CMV-FAM/TAMRA (Table 2). Cellular albumin (*ALB*) was amplified using primers 5’Alb and 3’Alb together with the hydrolysis probe Alb-FAM/Tamra (Table 2). The 20 µl PCR reactions contained 5 μl sample DNA together with 10 μl 2x SsoAdvanced Universal Probes Supermix (Biorad), 1 μl of each primer (5 μM stock solution), 0.3 μl probe (10 μM stock solution), and 2.7 μl H_2_O. For determination of reference C_T_ (cycle threshold) values, serial dilutions of the respective standards (10^8^-10^2^ DNA molecules of HCMV UL123 or *ALB*) were examined by PCR reactions in parallel. The thermal cycling conditions consisted of an initial step of 3 min at 95 °C followed by 40 amplification cycles (10 s at 95 °C, 30 s 60 °C). Viral genome copy numbers were subsequently calculated with the sample-specific C_T_ value set in relation to the standard serial dilutions.

### Growth analysis of HSV-1-GFP

Inducible HFF cells were seeded in triplicates into 96-well plates at a density of 1 × 10^4^ cells/well and were treated with doxycycline for 24 h, before they were infected with HSV-1 encoding a GFP-tagged VP22 a MOI of 0.1. At different times after infection, the GFP signal was measured in the living cells using a luminometer (SpectraMax iD5 Multi-Mode Microplate Reader, Molecular Devices) and the corresponding SoftMax Pro software.

### Quantification of RNA levels by reverse transcription real-time PCR (RT-qPCR)

For RNA isolation, 3 × 10^5^ to 6 × 10^5^ HFFs seeded in 6-well plates were lysed in 0.7 ml TRIzol (Invitrogen) for 5 min at RT. Total RNA was isolated using the Direct-zol Miniprep (Zymo Research) according to the manufacturer’s instructions, including an extended on-column DNase digestion for 30 min. Afterwards, first-strand cDNAs were synthesized from 0.3-0.5 μg of total RNA using oligo(dT)18 and random hexamer primers provided by the Maxima First Strand cDNA synthesis kit (Thermo Scientific). First-strand cDNAs were diluted 5-fold with nuclease-free water and subjected to quantitative SYBR Green PCR using an Agilent AriaMx Real-time PCR System with the corresponding software AriaMx 1.5 (Agilent Technologies, Inc.). Each reaction comprised 2 µl of sample or water along with 10 µl SsoAdvanced Universal SYBR Green Supermix (Biorad), 0.06 μl primer mix (50 μM stock solution), and 7.94 μl nuclease-free H_2_O. Utilized primers are listed in table 2. The initial denaturation step (95°C, 30 sec) was followed by 40 cycles of denaturation (95°C, 15 sec), primer annealing and strand elongation (60°C, 35 sec), followed by a dissociation stage to ascertain the specificity of the reactions. Data were analyzed using the 2^− ΔΔCt^ method with *GAPDH* (glyceraldehyde-3-phosphate dehydrogenase) as reference gene.

### Single-cell library preparation and sequencing

For single cell RNA sequencing, confluent HFF cells were infected with HCMV in T175 cm2 cell culture flasks. 6 hpi, cells were harvested and washed following instructions of the scRNAseq 10x Genomics Sample Preparation Demonstrated Protocol. From the cell suspension, 10000 live cells were subjected to single cell library preparation with a target cell count of 6000. Libraries were generated with the 10x Chromium Single Cell 3’ Gene Expression v3 kit (10x Genomics) according to the manufacturer’s instructions. Sequencing was performed on an Illumina HiSeq 2500 sequencer to a depth of about 140 million reads with a mean number of reads per cell greater than 20000. Reads were converted to FASTQ format using mkfastq from Cell Ranger v3.1.0 (10x Genomics) using the recommended read lengths for gene expression libraries. The gene reference data was constructed from the Cell Ranger human GRCh38 v3.0.0 reference by adding the HCMV reference genome (NCBI GeneBank id: BK000394.5). The BED format gene annotations were converted to GTF format using the bedToGenePred script from the UCSC Genome Browser utilities (http://hgdownload.cse.ucsc.edu/admin/exe/linux.x86_64/, accessed April 2020). Viral genome sequences and annotations were appended to the human GRCh38 sequences and annotations respectively. The reference data was constructed from these files using the Cell Ranger v3.1.0 mkref command with default settings. Reads were aligned to the custom reference using the Cell Ranger v3.1.0 count command. Human genes with less than 10 counts across all cells were excluded while HCMV genes were all included independent of expression level. The resulting cell data was filtered in Seurat v3.1.4 under R v3.6.3 (R Core Team 2020) to only retain cells with between 3000 and 28000 unique molecule identifier (UMI), more than 1500 genes, less than 10% mitochondrial, and more than 5% ribosomal reads per cell [54]. Applying these filters yielded 6462 high-quality cells. Gene expression values were normalized using Seurat LogNormalize, the top 2000 most variable features were selected using the vst method and then scaled to unit variance. Cell cycle states were scored using the CellCycleScoring function with the gene signatures provided in Seurat. UMAP embeddings were calculated using the RunUMAP function based on the top 20 PCA dimensions (n.neighbors = 10, min.dist = 0.3). Unsupervised clustering was performed using the Leiden algorithm on the two UMAP dimensions with a resolution of 0.1. The subclusters of the two larger communities visible in UMAP were manually merged into clusters 1 and 2 respectively, based on biological similarities. Differentially expressed genes were obtained using the MAST algorithm as implemented in the Seurat FindMarkers function [55].

### RNA sequencing

For RNA sequencing, HFF cells seeded in 6-well plates (3 × 10^5^ cells/well) were lysed in 0.7 ml TRIzol (Invitrogen) for 5 min at RT. Two wells were combined to obtainabout 6 × 10^5^ HFFs per sample. Total RNA was isolated using the Direct-zol Miniprep (Zymo Research) according to the manufacturer’s instructions, including an extended on-column DNase digestion for 30 min. Library preparation and sequencing was performed at Novogene (Cambridge, UK) using the Illumina NovaSeq 6000 sequencing platform to generate 2×150 bp paired-end libraries. The RNA-seq data were processed using the GRAND-SLAM pipeline (v2.0.7b) [37]. Adapter sequences were trimmed with Cutadapt (v3.5) using -a AGATCGGAAGAGCACACGTCTGAACTCCAGTCA -A AGATCGGAAGAGCGTCGTGTAGGGAAAGAGTGT. Trimmed reads were first aligned to rRNA (U13369.1) and Mycoplasma databases using Bowtie2 (v2.3.0) with default settings. Remaining reads were then mapped to a combined human and HCMV genome using STAR (v2.7.10b) with the parameter--alignEndsType Extend5pOfReads12. Genome references included human (Ensembl, release 90) and HCMV (GenBank: X17403). Following alignment, reads were converted to CIT files using GEDI Bam2CIT, corrected with GEDI CorrectCIT, and merged with GEDI MergeCIT. These CIT files were then processed with GRAND-SLAM (-trim5p 15-modelall-intronic) to generate gene-level read counts. Downstream analyses were conducted using the grandR package (v0.2.1) [56, 57]. Filtering and normalization were performed twice: initially with NormalizeTPM() and FilterGenes() retaining only genes with TPM ≥ 30 in at least one condition and subsequently filtered with FilterGenes() to retain genes with sufficient expression and normalized using Normalize(). Differential expression was assessed using Pairwise() to compare ISG20-overexpressing samples with controls. Genes with FDR q <0.05 and |Log2(fold change)| >1 were considered significant. The Enrichr Web Server (https://maayanlab.cloud/Enrichr/) was utilized to perform Reactome enrichment analysis of differentially expressed genes (DEGs) [58]. TEs were quantified using the snakePipes <u>ncRNAseq</u> pipeline starting from the aligned reads [59]. This pipeline quantifies TEs using TEtranscripts and then identifies differentially expressed TEs using DESeq2 [60, 61].

### SLAM-seq analysis

To label newly synthesized DNA, HFF cells (3 × 10^5^ cells/well) were treated with 4-thiouridine (4sU) at a final concentration of 400µM for 1 h prior to cell lysis. Cells were lysed TRIzol for 5 min at room temperature and kept at −80°C. Two wells were combined to obtain around 6 × 10^5^ cells per sample. Total RNA was isolated using the Direct-zol Miniprep kit (Zymo Research) according to the manufacturer’s instructions, including an extended on-column DNase digestion for 30 min. The 4sU alkylation reaction was performed by incubating 2 to 5 μg total RNA in 1× RNase-free PBS (pH 8) containing 50% dimethyl sulfoxide (DMSO) and 10 mM iodoacetamide (IAA) at 50°C for 15 min. The reaction was quenched with 100 mM DTT and incubated for 5 min on ice, before the RNA was purified using the RNeasy MinElute Cleanup Kit (Qiagen) according to the manufacturer’s instructions. Library preparation and sequencing was performed at Novogene (Cambridge, UK) using a Illumina NovaSeq X Plus sequencing platform to generate 2×150 bp paired-end libraries. The SLAM-seq data were processed using the GRAND-SLAM pipeline (v2.0.7b) [37]. Reads were aligned to rRNA (U13369.1) and Mycoplasma databases with Bowtie2 (v2.3.0), followed by alignment to a combined human and HCMV genome using STAR (v2.7.10b) with the following parameters:--outFilterMismatchNmax 20,--outFilterScoreMinOverLread 0.4, -- outFilterMatchNminOverLread 0.4, and --alignEndsType Extend5pOfReads12. Aligned reads were converted, corrected, and merged using GEDI (Bam2CIT, CorrectCIT, and MergeCIT) [62]. GRAND-SLAM was then used (-trim5p 15-modelall-intronic) to generate read counts and new-to-total ratios (NTRs) at the gene level. Downstream analyses were performed using the **grandR** package (v0.2.5) [56, 57]. Genes were filtered with FilterGenes() and normalized with Normalize(). 4sU dropout was evaluated using Plot4sUDropoutAll(), and half-lives were estimated with FitKineticsSnapshot(). Upregulated genes (LFC >1, q <0.05) were identified using GetSignificantGenes() and then filtered to include only genes with the ’ZNF’ prefix. Promoter regions (1 kb upstream, 100 bp downstream of TSS) were extracted using promoters() from GenomicRanges with gene coordinates from TxDb.Hsapiens.UCSC.hg38.knownGene. Gene mappings were obtained via org.Hs.eg.db, and sequences were retrieved from BSgenome.Hsapiens.UCSC.hg38, excluding _alt annotations. Motif analysis was performed with monaLisa (v1.0.0) using JASPAR2020 position frequency matrices (PFMs) [63]. PFMs were converted to position weight matrices (PWMs) with TFBSTools, and promoter sequences of upregulated ZNF genes were compared against a random genomic background. Motifs with negLog10Padj >4 were considered significant.

## Author Contributions

Data curation: MH, MS, PK, TR, FE, CCF

Conceptualization: TS, MS, FE

Formal analysis: MH, MS, CCF, FE

Funding acquisition: TS, CCF, FE

Investigation: MH, MS, E-MR, CP, TR, PK, NK, AR, AKK, ACK

Methodology: CP, TR, PK, NK, AR, AKK, ACK, CCF, FE

Project administration: TS

Supervision: TS, MS

Visualization: MH, MS, TR, CCF, FE, TS

Validation: MH, MS

Writing – original draft: TS, MS

Writing – review & editing: TS, MS, CCF, FE

