## Supplementary Figures for "Exonuclease ISG20 inhibits human cytomegalovirus replication by inducing an innate immune defense signature"

### Supplementary Figures; Hehl et al.

#### Supplementary Figure 1:

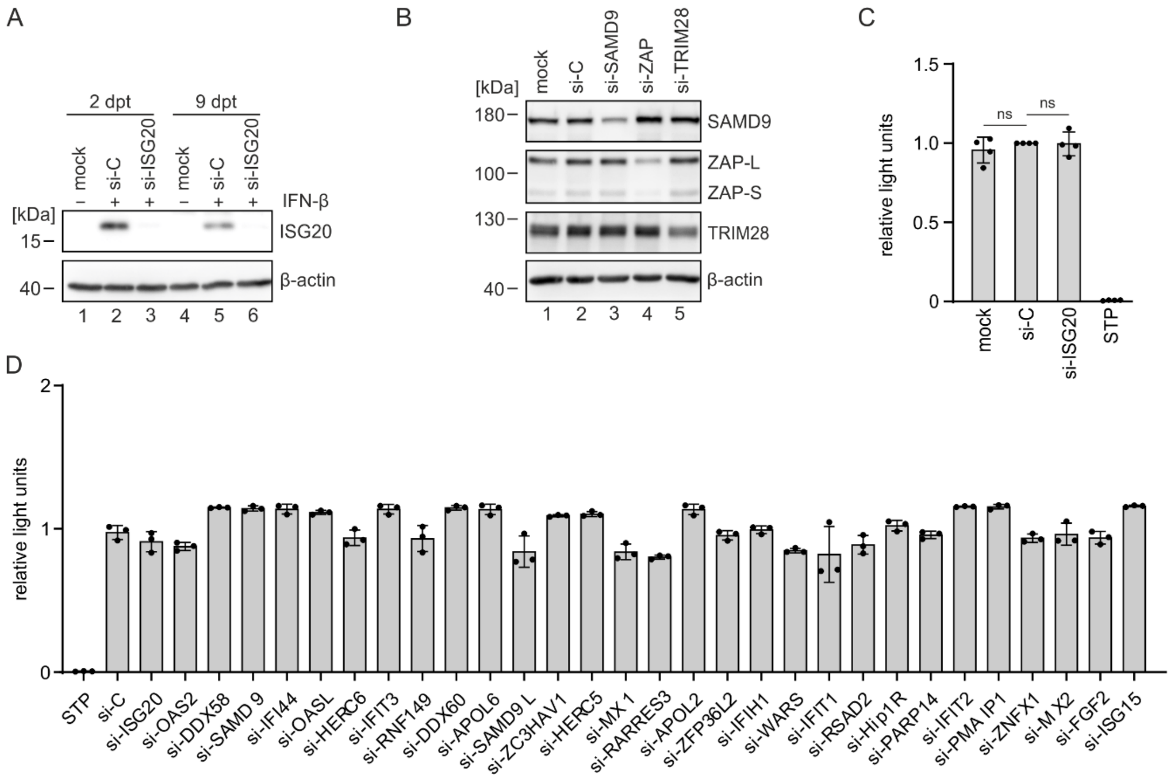


**Supplementary Figure 1**. **Effects of siRNA transfection on cell viability and expression of target genes**. (A) Knockdown efficiency of ISG20. HFF cells were not treated (mock), transfected with control siRNA (si-C) or with ISG20-specific siRNA (si-ISG20) at a final concentration of 28.3 nM. To enable the detection of ISG20 by western blotting, cells were treated with IFN-β (1000 U/ml) for 24 h, before they were harvested at 2 and 9 after transfection (dpt) corresponding to the time point of infection and measurement in Figure 1D, respectively. Β-actin was included as loading control. (B) Knockdown efficiency of proteins included in the screening in Figure 1D. HFF cells were not treated (mock), transfected with control siRNA (si-C) or siRNAs specific for SAMD9, ZAP and TRIM28 at a final concentration of 28.3 nM. 2 days after transfection, cells were harvested for western blot analysis of SAMD9, ZAP and TRIM28 protein levels. Β-actin was included as loading control. (C, D) Analysis of cell viability after siRNA transfection of HFF cells. HFFs were seeded in triplicates and, one day later, transfected with control siRNA (si-C) or siRNA against indicated proteins at a final concentration of 28.3 nM, or were left untreated (mock). 9 days after transfection, corresponding to the measurement time point in Figure 1D, intracellular ATP levels were measured using the CellTiterGlo assay. The cytotoxic kinase-inhibitor staurosporin (STP) was used as control and was added 24 h before the measurement at a concentration of 5 µM. Depicted are mean values +/-SD of four independent experiments (C) or triplicates (D). Statistical analysis was performed using a one-sample t-test. Ns, not significant. dpt, days post-transfection.

#### Supplementary Figure 2:

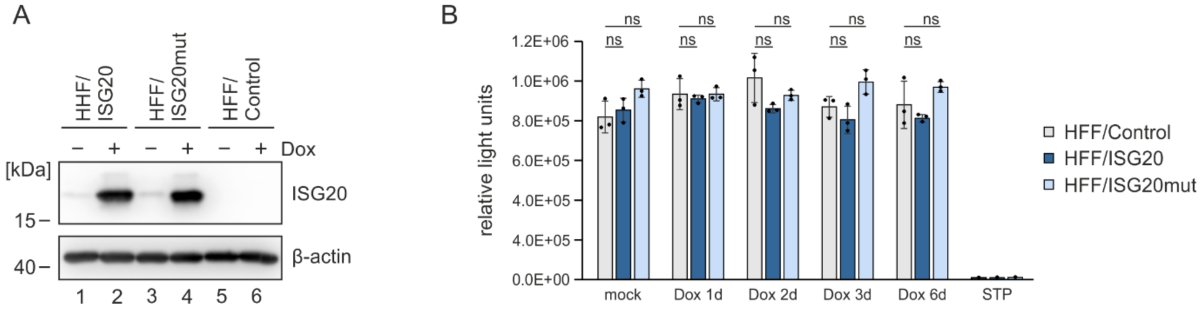


**Supplementary Figure 2. Generation of HFF with inducible expression of ISG20 or ISG20mut.** (A) Induction of ISG20 expression with doxycycline. HFF/Control, HFF/ISG20 and HFF/ISG20mut were stimulated with 500 ng/ml doxycycline (Dox) for 24 h and ISG20 expression was analyzed by western blotting. Cellular β-actin levels served as loading control. (B) Analysis of cell viability in HFF with doxycycline-inducible expression of ISG20. HFF/Control, HFF/ISG20 and HFF/ISG20mut were seeded in triplicates and either not induced (mock), induced with 500 ng/ml doxycycline (Dox) for indicated times (1 to 6 days) or treated with the cytotoxic kinase-inhibitor staurosporin (STP) at a concentration of 5 µM for 1d. Cell viability was determined by measuring intracellular ATP levels using the CellTiter-Glo Assay. Values are shown as means +/- SD of triplicate samples. One representative experiment out of two is shown. Dox, docycycline.

#### Supplementary Figure 3

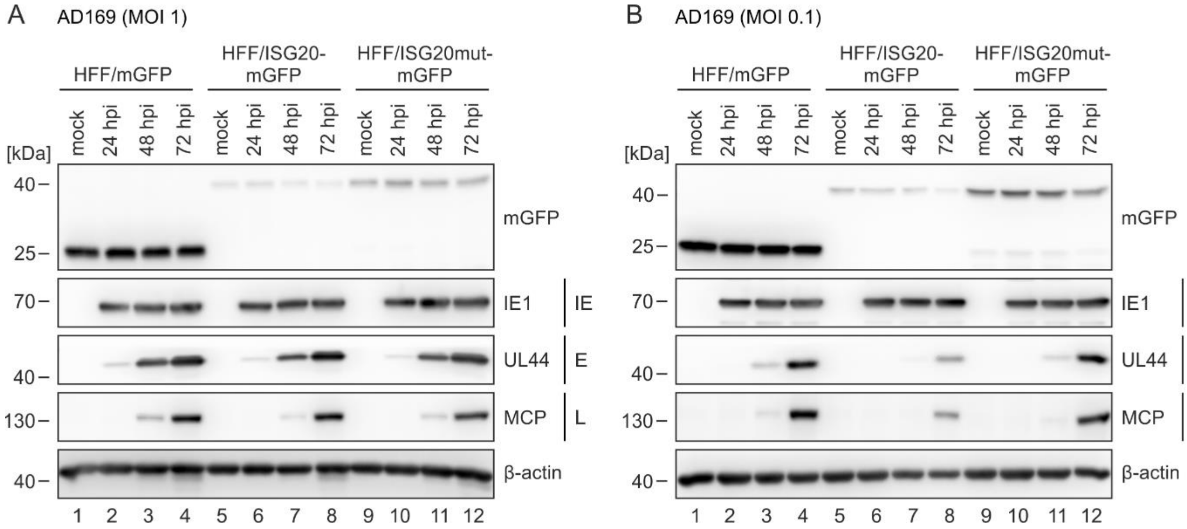


**Supplementary Figure 3. Effect of ISG20-mGFP on the expression pattern of HCMV proteins**. HFF/mGFP (control cells), HFF/ISG20-mGFP and HFF/ISG20mut-mGFP were not infected (mock) or infected with HCMV strain AD169 a MOI of 1 (A) or 0.1 (B). At indicated times after virus inoculation, cell lysates were prepared and the expression levels of viral immediate early (IE), early (E), and late (L) proteins were analyzed by western blotting. Cellular β-actin levels served as loading control. One representative experiment of two is shown.

#### Supplementary Figure 4

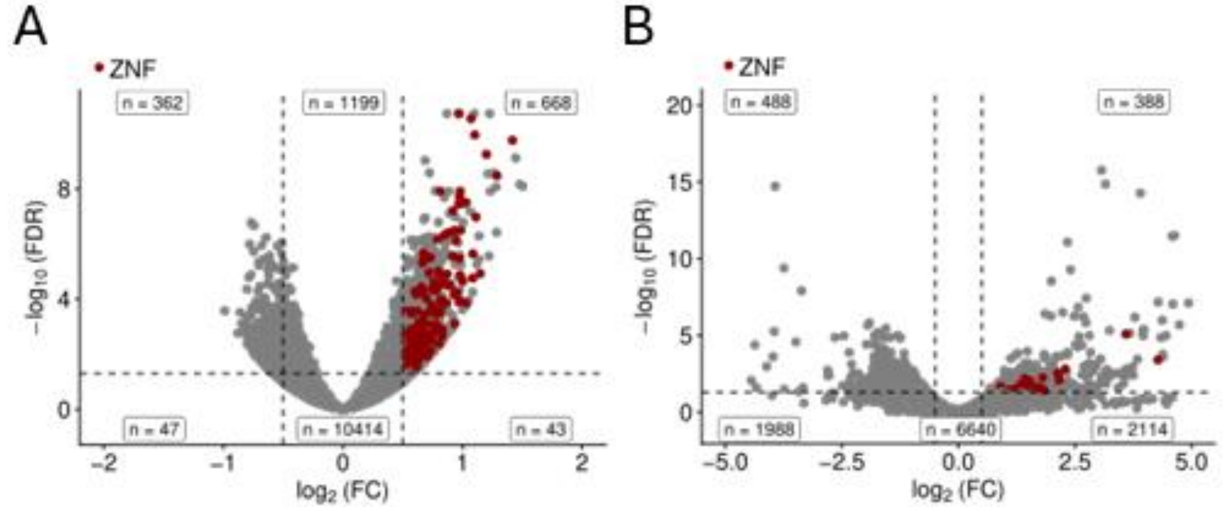


**Supplementary Figure 4. Effect of ISG20 on the cellular transcriptome.** Volcano plots showing differentially expressed genes in HEK293T (A) and MEF (B) cells. Colored dots represent significantly regulated ZNF genes with adjusted p-value <0.05 and |log2 fold change) >0.5|.
